# Pharmacological PINK1 activation ameliorates Pathology in Parkinson’s Disease models

**DOI:** 10.1101/2023.02.14.528378

**Authors:** Randall M. Chin, Rishi Rakhit, Dara Ditsworth, Chengzhong Wang, Johan Bartholomeus, Song Liu, Akash Mody, Alex Laishu, Andrea Eastes, Chao Tai, Roy Y. Kim, Jessica Li, Steven Hansberry, Saurabh Khasnavis, Victoria Rafalski, Donald Herendeen, Virginia Garda, Jennie Phung, Daniel de Roulet, Alban Ordureau, J. Wade Harper, Shawn Johnstone, Jan Stöhr, Nicholas T. Hertz

## Abstract

PINK1 loss-of-function mutations and exposure to mitochondrial toxins are causative for Parkinson’s disease (PD) and Parkinsonism, respectively. We demonstrate that pathological α-synuclein deposition, the hallmark pathology of idiopathic PD, induces mitochondrial dysfunction and impairs mitophagy, driving accumulation of the PINK1 substrate pS65-Ubiquitin (pUb) in primary neurons and in vivo. We synthesized MTK458, a brain penetrant small molecule that binds to PINK1 and stabilizes an active heterocomplex, thereby increasing mitophagy. MTK458 mediates clearance of α-synuclein pathology in PFF seeding models in vitro and in vivo and reduces pUb. We developed an ultrasensitive assay to quantify pUb levels in plasma and observed an increase in pUb in PD subjects that correlates with disease progression, paralleling our observations in PD models. Our combined findings from preclinical PD models and patient biofluids suggest that pharmacological activation of PINK1 is worthy of further study as a therapeutic strategy for disease modification in PD.

**Highlights:**

1. Discovery of a plasma Parkinson’s Disease biomarker candidate, pS65-Ubiquitin (pUb)
2. Plasma pUb levels correlate with disease status and progression in PD patients.
3. Identification of a potent, brain penetrant PINK1 activator, MTK458
4. MTK458 selectively activates PINK1 by stimulating dimerization and stabilization of the PINK1/TOM complex
5. MTK458 drives clearance of α-synuclein pathology and normalizes pUb in in vivo Parkinson’s models

## Introduction

Parkinson’s Disease (PD) is the second most common neurodegenerative disease, with more than 10 million patients affected worldwide (Ball et al., 2019) and no FDA approved disease modifying therapies (Bloem et al., 2021; Kim et al., 2018; Yang et al., 2020). Hallmark pathologies of idiopathic PD include the formation of Lewy bodies consisting primarily of aggregated α-synuclein and ubiquitin, neuronal loss in the substantia nigra, striatal dopamine deficiency, and mitochondrial deficits (Poewe et al., 2017). These neuropathologic features result in motor deficits such as tremors, bradykinesia, stiffness, and impaired balance, as well as non-motor symptoms such as cognitive impairment. The foundation of therapeutic treatment for PD consists of dopamine replacement *(e.g.,* Levodopa), which loses clinical efficacy and, with increased disease duration, can cause dyskinesia and dystonia (Fox et al., 2018; Pringsheim et al., 2021), and has no effect on the underlying pathologic progression of the disease. No disease modifying therapies are available, although many approaches are in preclinical and clinical development (Lang et al., 2022; Pagano et al., 2022).

Further complicating the development of disease modifying therapeutics is the absence of biomarkers to monitor disease progression at a molecular level (Parnetti et al., 2019). Unlike the multitude of disease progression biomarkers and PET ligands available for Alzheimer’s Disease (AD), there are no approved physiologic, radiologic, or blood-based biomarkers to confirm the clinical diagnosis or monitor progression of PD (Parnetti et al., 2019). One assay to detect aggregated α-synuclein, the real-time quaking-induced conversion (RT-QuIC) can identify PD patients from cerebrospinal fluid (CSF), but remains a binary disease readout (Brockmann et al., 2021; Fairfoul et al., 2016; Poggiolini et al., 2022). While large-scale efforts are underway to identify disease progression biomarkers, including the Michael J. Fox Foundation (MJFF)-sponsored LRRK2 Cohort Consortium and Parkinson’s Progression Markers Initiative <u>(MJFF PPMI)</u>, these efforts have not yet identified an ideal biomarker (Parnetti et al., 2019).

Recent advances in the understanding of key biological mechanisms leading to PD and/or affecting disease progression are helping to inform a new generation of biomarkers and therapeutics. Various familial forms of PD have been identified and implicate α-synuclein aggregation, lysosomal dysfunction, and notably, impairments in mitochondrial quality control mechanisms as causative for Parkinson’s disease (Borsche et al., 2021). Homozygous recessive mutations in Parkin, a mitochondrial associated E3-Ubiquitin ligase, or in PINK1, a serine/threonine kinase that serves as a master regulator of mitochondrial quality control, cause DA neuron degeneration and early-onset PD (Kitada et al., 1998; Valente et al., 2004)

Beyond patient genetics, multiple lines of experimental evidence suggest that PD is intimately linked to mitochondrial function, so it has been proposed that improving mitochondrial quality control and homeostasis could be a viable approach for disease modification in PD. It is well established that nigrostriatal dopaminergic (DA) neurons, which degenerate in PD, are particularly susceptible to mitochondrial dysfunction (Fang et al., 2019; Henchcliffe and Beal, 2008). Multiple mitochondrial toxins, including MPTP, rotenone, and paraquat, are linked to Parkinsonism in humans and have been shown to drive preferential degeneration of dopaminergic neurons in in vivo models of PD (Nandipati and Litvan, 2016). Additionally, α-synuclein aggregates have been shown to accumulate at mitochondria, interfere with mitochondrial function, and delay mitophagy (Devi et al., 2008; Ryan et al., 2018; Shaltouki et al., 2018; Wang et al., 2019). Intriguingly, an electron microscopy study of PD patients’ brains identified mitochondrial components within Lewy Bodies (Shahmoradian et al., 2019), further highlighting a potential mitochondria-pathogenic α-synuclein connection. Despite extensive evidence, the hypothesis that ameliorating mitochondrial function may be beneficial for PD has not been thoroughly tested in humans; the clinical trials run to date have focused on using antioxidants to prevent damage rather than augmenting mitochondrial quality control pathways to address the underlying dysfunction (Borsche et al., 2021; Kim et al., 2021). In this work, we explore the idea that improving mitochondrial function by activating PINK1 may be beneficial for PD.

Depolarization-induced mitochondrial autophagy (mitophagy) is tightly regulated by PINK1/Parkin pathway activation. Under basal conditions, PINK1 is targeted to the mitochondrion, cleaved by PARL, and exported for degradation by the proteosome, preventing mitochondrion-localized kinase activity. (Deas et al., 2011; Greene et al., 2012; Jin et al., 2010; Lazarou et al., 2012; Yamano and Youle, 2013). Conversely, in cells undergoing mitochondrial stress, the full length 63 kDa PINK1 is stabilized on the outer mitochondrial membrane and becomes catalytically active. Active PINK1 forms a high molecular weight complex with TOM where it dimerizes and autophosphorylates at Serine 228, potentiating PINK1 activation (Kakade et al., 2022; Lazarou et al., 2013; Okatsu et al., 2012; Ordureau et al., 2014). Active PINK1 directly phosphorylates ubiquitin (Ub) and the ubiquitin-like domain of Parkin, at the homologous Serine residue (Kane et al., 2014; Kazlauskaite et al., 2014; Kondapalli et al., 2012; Koyano et al., 2014; Ordureau et al., 2014; Wauer et al., 2015). Both PINK1-mediated phosphorylation of Parkin and binding to pUb drive activation of Parkin, ultimately leading to the recruitment of additional Parkin to the mitochondrion. Parkin ubiquitinates outer mitochondrial membrane proteins such as the mitofusins (Antico et al., 2021; Bingol et al., 2014; Ordureau et al., 2020; Sarraf et al., 2013), leading to their degradation and causing fragmentation of the stressed parts of the mitochondrial network. The damaged and fragmented pieces of mitochondria are subsequently engulfed by the autophagosome in the process of mitophagy. Mitophagy is believed to be the primary means of disposing of damaged mitochondria, however, if the mitophagy machinery is compromised or insufficient, cells can release their damaged mitochondria extracellularly (Choong et al., 2021; Tan et al., 2022).

Since mitochondrial dysfunction is a key element of PD and LoF mutations in PINK1 result in PD, we sought to develop small molecule activators of PINK1 as a treatment for PD. Building upon our previous discovery of a novel role for kinetin as an activator of PINK1 (Hertz et al., 2013), we synthesized and tested kinetin analogs seeking compounds with superior pharmacological properties leading to the discovery of MTK458. To further understand the role of mitochondrial dysfunction in PD, we took advantage of the fact that pUb is a PINK1-specific metabolite and a mitochondrial damage indicator to measure pUb levels in the plasma of PD patients. We discovered that plasma pUb levels are significantly increased in the PD population compared to healthy controls, indicating an increase in mitochondrial dysfunction with PD. We further demonstrated that cells or mice challenged with α-synuclein aggregation exhibit increased pUb, mitochondrial dysfunction, and stalled mitophagy, consistent with our human plasma pUb data. Remarkably, MTK458 dosing can both alleviate mitochondrial stress, as evidenced by a decrease in plasma pUb, and decrease the levels of α-synuclein aggregation. These data suggest that pharmacological agonists of PINK1 have the potential to treat patients by addressing the pathological hallmarks of PD.

## Results

### Pre-formed fibrils of α-synuclein cause mitochondrial dysfunction and impair mitophagy

The pathological hallmark of idiopathic and most genetic forms of PD is the misfolding and subsequent deposition of α-synuclein into insoluble, beta sheet rich aggregates deposits referred to as Lewy pathology, and can be detected by phosphorylation of α-synuclein at Serine 129 (p129) (Rocha et al., 2018). High-resolution studies revealed that mitochondrial fragments are an integral part of Lewy Body structures, suggesting an interaction of misfolded α-synuclein with mitochondrial structures (Shahmoradian et al., 2019). Furthermore, patient-derived A53T SNCA iPSCs show delayed mitophagy (Devi et al., 2008; Ryan et al., 2018; Wang et al., 2019). Consistent with these findings, we found pS129 α-synuclein associated nearly exclusively with the mitochondrial fraction of cortical brain extracts in human PD patients (Figure 1A-C). To assess whether pathogenic α-synuclein induced mitochondrial dysfunction we seeded cultured primary neurons with α-synuclein preformed fibrils (PFFs), which lead to pS129 α-synuclein aggregation (Figures 1D-E and S1A-B) (Volpicelli-Daley et al., 2011). We observed concentration- and time-dependent defects in mitochondrial respiration (Figures 1F-I and S1C-F), impaired mitophagy (Figures 1J), and a dose-dependent accumulation of pUb (Figure 1K). Interestingly, a chronic, low dose of the mitochondrial uncoupler CCCP also led to impaired mitophagy in neurons (Figure 1J). This result suggests that stimulating chronic low levels of mitophagy via mitochondrial uncoupling leads to stalled mitophagy in which PINK1 is partially activated but unable to complete the mitophagic process; as a result, pUb that would otherwise be turned over when the mitophagic process is completed builds up in the cell. These data support a model in which α-synuclein pathology increases mitochondrial dysfunction and impairs mitophagy, leading to accumulation of pUb (Figure 1L).

**Figure 1:**
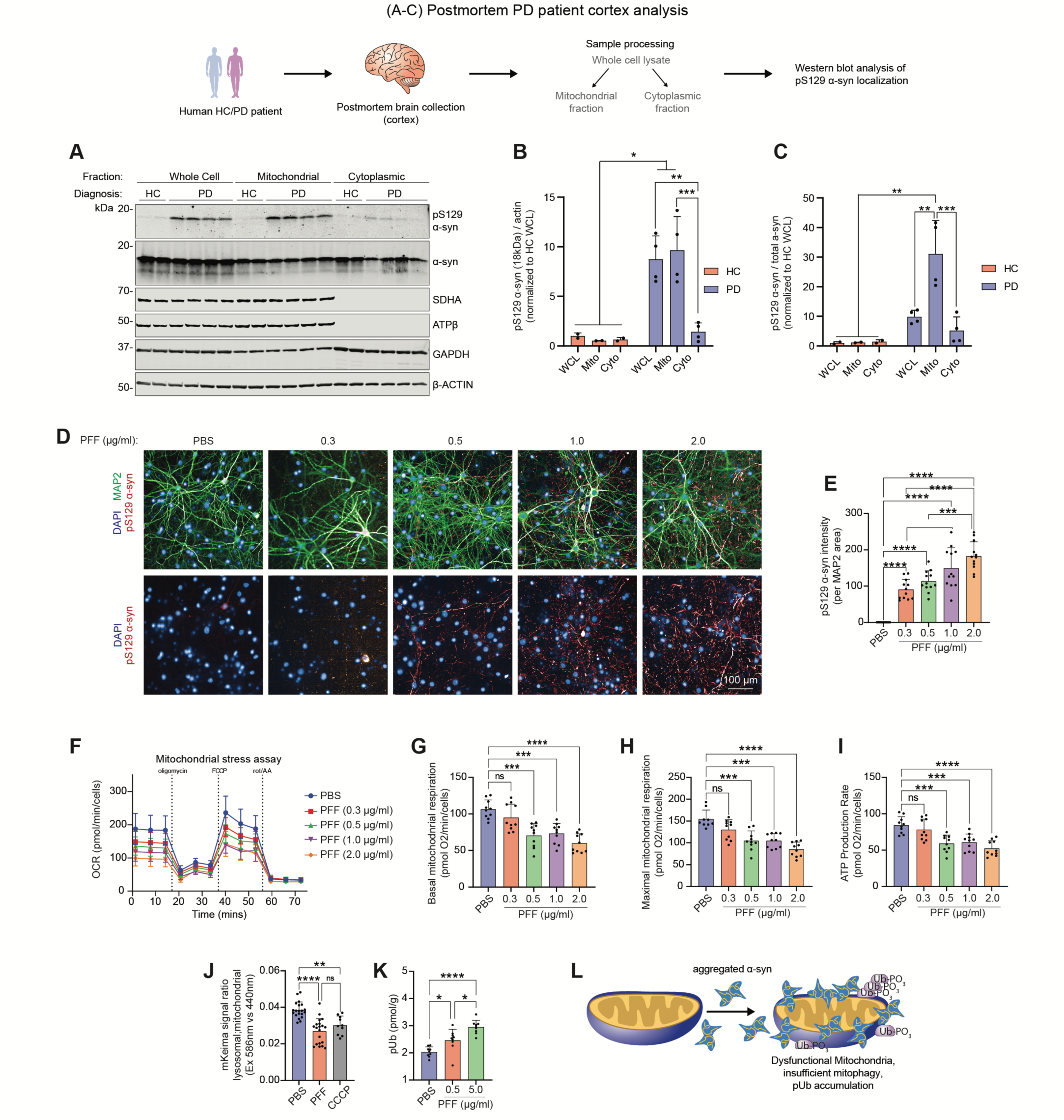
Mitochondrial associated α-synuclein is present in PD patient brain tissue and induces mitochondrial dysfunction in primary neurons. (A) Human brain pieces from HC or PD individuals were separated into mitochondrial or cytoplasmic fractions, or unfractionated as whole cell lysate, and analyzed by immunoblotting. pS129 α-syn is present in the mitochondrial fraction. (B-C) Quantification of (A) is shown. (D-J) M83 primary neurons were challenged with various concentrations of PFFs, and after 21 days, pS129 α-synuclein staining (D-E), mitochondrial respiration (F-I), and mitophagy (J) was assessed. (D-E) pS129 α-synuclein intensity is increased dose dependently with PFF challenge. (F-I) PFF challenge dose dependently decreases basal respiration, maximal respiration, and ATP production rate. (J) Primary neurons challenged with 21 days of 1 µg/ml PFFs or 10 nM CCCP show impaired mitophagy as measured by the mKeima reporter. (K) Primary neurons challenged with mPFFs show an accumulation in pUb as measured by the MSD pUb assay. (L) A model of our hypothesis: aggregation of pS129 α-syn occurs at the mitochondria, leading to dysfunctional mitochondria, insufficient mitophagy, and accumulation of pUb. Unless otherwise indicated, mean and SD is shown; one-way ANOVA was used for statistical analysis. Mean ± SD. *p < 0.05, **p < 0.01, ***p < 0.001, ****p < 0.0001 n.s., not significant. trend where pUb levels increase with disease severity. (J) Patient plasma pUb levels are binned according to their gender and LRRK2 genotype. The increase in plasma pUb in the PD population occurs independently of gender or LRRK2 mutation. Unless otherwise noted, mean and SD are shown; one-way ANOVA or t-test was used for statistical analysis. *p<0.05, **p<0.01, ***p<0.001, ****p<0.0001, n.s., not significant.

### Parkinson’s disease patients have elevated levels of PINK1 substrate pUb in plasma

Aggregated α-synuclein represents an underlying pathobiology of PD and induces mitochondrial dysfunction in primary neurons as evidenced by an increase in pUb (Figure 1K). A recent publication showed a significant increase in the PINK1 substrate pUb in postmortem brain tissue taken from PD patients (Hou et al., 2018), substantiating an increase in mitochondrial dysfunction in the central nervous system (CNS) of PD patients; however, there are currently no published reports of pUb measurement in a peripherally accessible biofluid. Furthermore, it has been shown that cells can release their damaged mitochondrion (containing pUb) extracellularly, which provides neurons an alternative way to clear mitochondria (Choong et al., 2021; Tan et al., 2022). Given that pUb had been shown to be elevated in PD brain tissue and that cells can release damaged mitochondria, we hypothesized that brain-derived pUb would be detectable and potentially elevated in the biofluids of PD patients.

In order to quantify pUb in human plasma, we developed a robust, ultrasensitive plasma biomarker assay on the SMCxPro (previously Singulex) platform (Cohen and Walt, 2019) to measure pUb. PINK1 is the only kinase known to phosphorylate Ubiquitin at Serine 65 (Kane et al., 2014; Kazlauskaite et al., 2014; Koyano et al., 2014; Ordureau et al., 2014). Samples from PINK1 knockout (KO) animals analyzed by the SMCxPro assay showed significantly lower levels of pUb, demonstrating the specificity of the assay (Figure S2A-D). The SMCxPro pUb assay also passed our assay validation criteria for parallelism, dilution linearity, spike recovery, test/re-test reliability, and precision (Figures S2E-I).

Having established the assay, we analyzed >1600 blinded plasma samples from two independent cohorts within the MJFF LRRK2 consortium cohort (LCC, Figures 2A-B and S2A). We found significantly elevated plasma pUb in each cohort (BioRep p<0.0001, Tel Aviv p<0.0001) (Figures 2B-D); combined, we found that plasma pUb levels are significantly increased by 22.4% in the plasma of PD patients vs healthy controls, ROC AUC=0.67, OR 2.62 (Figure 2E-F). There is a small increase in plasma pUb with age (Figure 2G); however, in comparing age-matched HC and PD patients, there is still a significant increase in plasma pUb in the PD cohort, indicating that the increase in plasma pUb in PD is independent of aging (Figure 2G). Remarkably, plasma pUb correlates with the UPDRS Part 3 and Modified H&Y clinical progression scores (Figures 2H-I). The increase in plasma pUb in the PD population occurs independently of gender, LRRK2 mutation status, levadopa usage, coffee consumption, or cigarette smoking (Figures 2J and S3B-E). These data suggest that pUb could represent an idiopathic PD progression biomarker, linking more common idiopathic PD with a rare genetic factor cause, PINK1.

**Figure 2:**
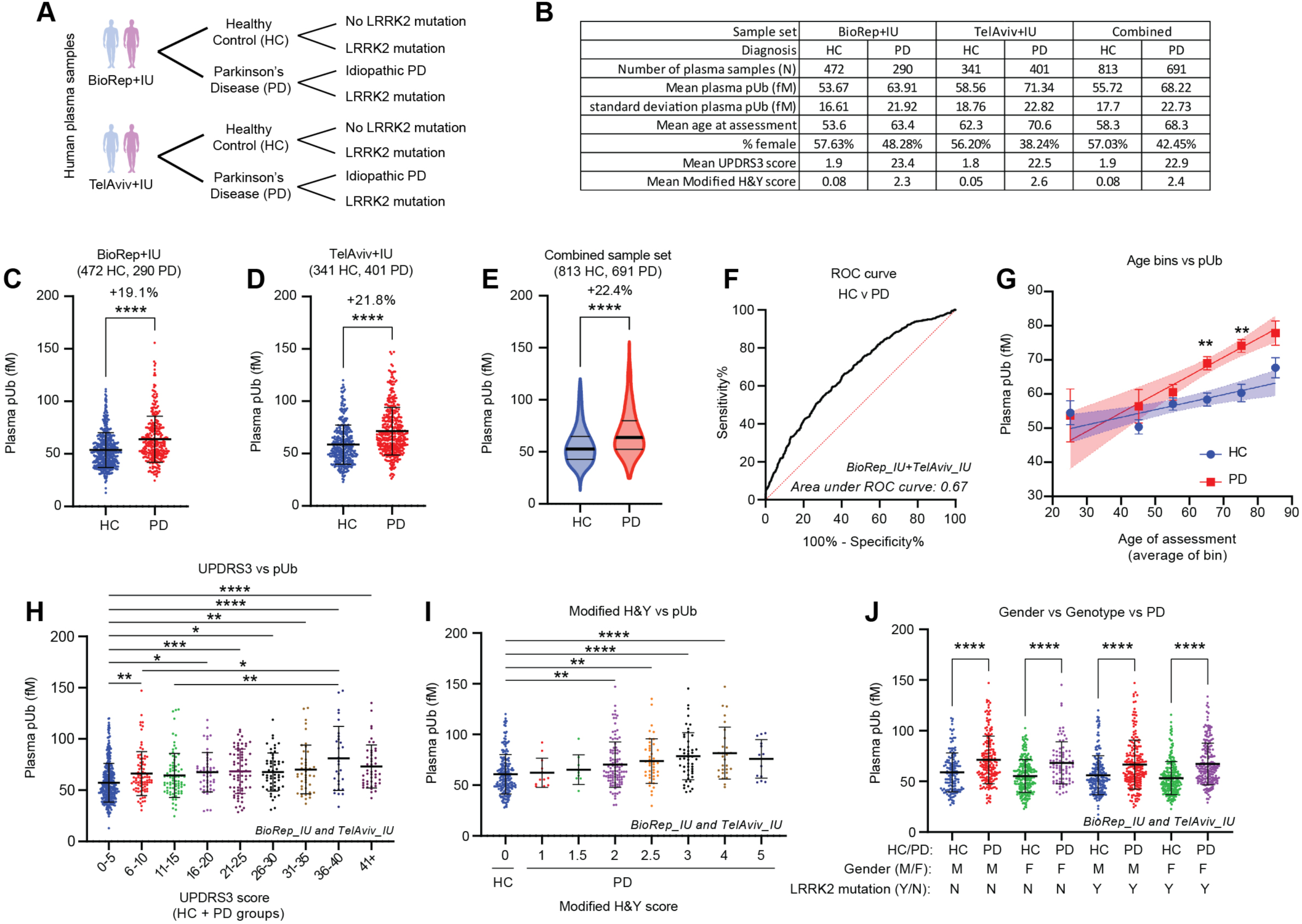
PINK1 substrate pS65-Ub increased in plasma as a general feature of idiopathic Parkinson’s Disease. (A) Human plasma samples were obtained from the BioRep+IU or TelAviv+IU cohorts from the MJFF LCC study. HC or PD patients, with or without LRRK2 mutations, were included. (B) Table describing the number of samples, mean age, gender, and clinical disease progression scores of each cohort. (C-E) Using the SMCxPRO pUb assay, plasma pUb levels were quantified in HC or PD patients from either cohort separately or combined. For all other figures with patient samples, we sourced data from the combined sample set. (F) An ROC curve based on Figure 1E is shown, with an area under the ROC curve of 0.67. (G) Patient plasma pUb levels are binned according to their age at assessment. The increased levels of plasma pUb in the PD population occurs independently of age. Mean and SEM are shown. (H) Patient plasma pUb levels are binned according to their total UPDRS Part 3 (UPDRS3) score, showing a trend where pUb increases with UPDRS3 score. (I) Patient plasma pUb levels are binned according to their Modified H&Y score, showing a trend where pUb levels increase with disease severity. (J) Patient plasma pUb levels are binned according to their gender and LRRK2 genotype. The increase in plasma pUb in the PD population occurs independently of gender or LRRK2 mutation. Unless otherwise noted, mean and SD are shown; one-way ANOVA or t-test was used for statistical analysis. *p<0.05, **p<0.01, ***p<0.001, ****p<0.0001, n.s., not significant.

The observation that pUb is elevated in PD patients suggests two possible hypotheses. One is that higher levels of PINK1 activity, as indicated by higher pUb, are causative for PD. This idea is flawed because PINK1 LoF mutations (reduced PINK1 activity) are genetically linked with PD. The other hypothesis is that pUb accumulation is a byproduct of PD, in line with the model and supportive data above (Figures 1J-K) that connect increased mitochondrial dysfunction and impaired mitophagy to an increase in pUb. In the second hypothesis, alleviation of mitochondrial stress and stalled mitophagy by further activation of PINK1 would result in lower pUb than the stressed condition. It would then be predicted that small molecule activators of PINK1 could provide therapeutic benefit in idiopathic PD, potentially leading to a disease modifying therapy. To test this concept, we sought to identify such molecules.

### Identifying and qualifying small molecule activators of PINK1

Our experiments establish that reduced rates of mitophagy and unresolved mitochondrial damage are key impairments in PD pathobiology and support pharmacological activation of PINK1 as a potential avenue for therapeutic intervention. Our original approach to activate PINK1 with neo-substrates (Hertz et al., 2013) led us to the discovery of kinetin, which activates PINK1 in cells and relieves mitochondrial mutations in flies and mice in a PINK1-dependent manner (Osgerby et al., 2017; Tsai et al., 2022). However, because kinetin has low potency and poor pharmacokinetics (PK) and brain penetrance, we could not detect an effect in PD models in vivo (Orr et al., 2017). To overcome these limitations and to discover novel small molecule PINK1 activators with drug-like properties, we synthesized and screened small molecule analogs of kinetin. Active compounds were selected by measuring activity in a cell-based assay for mitophagy in which a pH sensitive protein (keima) is localized on mitochondria (mKeima) and a characteristic shift in the absorption/excitation spectrum is observed upon initiation of mitophagy (Figure S4A-B) (Lazarou et al., 2015). We tested each compound in a 7-point concentration curve in the presence of a low concentration (1 μM) of FCCP and oligomycin (FO). In cell culture models, low levels of FO are necessary to trigger mitochondrial stress and stabilize PINK1; this dose was selected because it did not robustly trigger mitophagy on its own.

In order to rule out additive mitochondrial toxicity as the mechanism of action of a compound showing activity in the mKeima assay, we further counter-screened active compounds for mitochondrial toxicity in the galactose/glucose assay (Arroyo et al., 2016; Gohil et al., 2010; Marroquin et al., 2007). We used a 20% decrease in growth rate in galactose media relative to glucose media as a cutoff for mitotoxicity (Figures 3A-B and S4C). A subset of these active, non-mitotoxic compounds were then evaluated for initial developability using the following criteria: solubility, permeability, brain efflux, in vitro liver microsome clearance in multiple species, CYP screening, plasma protein binding, and hERG inhibition (Figure S4A). Several compounds that fulfilled developability criteria were tested in mouse pharmacokinetic (PK) and tissue distribution studies. The compound MTK458 showed good potency, no observable mitotoxicity, attractive oral PK, and high brain penetrance (Figure 3C, Data will be submitted upon full review). To further confirm that MTK458 is not a mitochondrial toxin, we measured mitochondrial respiration in HeLa cells treated with MTK458 for 1 hour. MTK458 did not affect basal respiration, maximal respiration, or spare respiratory capacity (Figure S4D).

**Figure 3:**
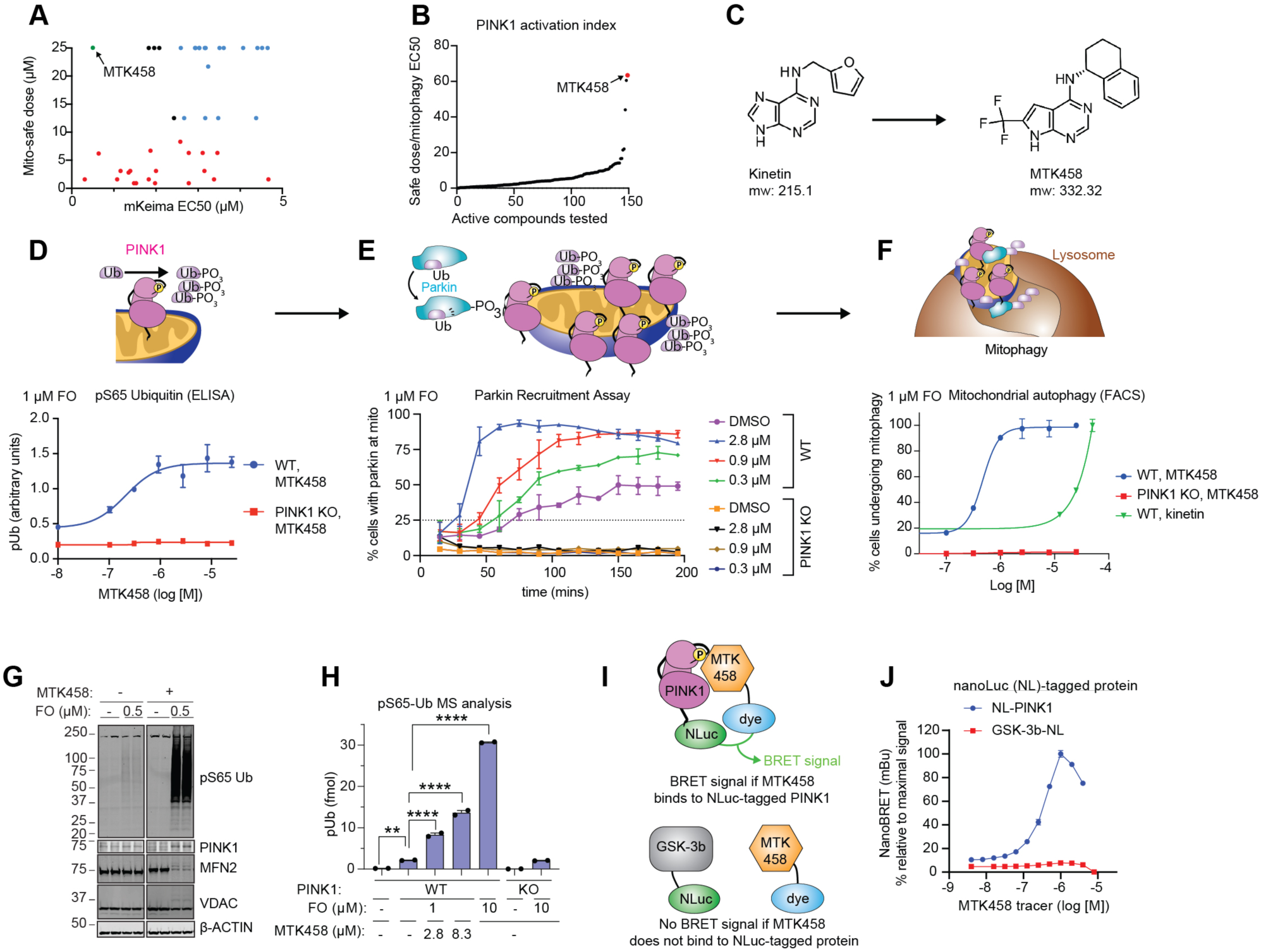
Identification of the small molecule PINK1 activator MTK458. (A-B) Compounds were screened for their ability to induce mitophagy in the mKeima assay and counter-screened for mitochondrial toxicity in the galactose/glucose assay. EC50 or EC20 measurements for either assay, respectively, are graphed. MTK458 is one of the best compounds in terms of high potency and low toxicity. (C) The structures of kinetin and MTK458 are shown. (D) YPMK (WT or PINK1 KO) cells were treated with a dose range of MTK458 and 1 µM FCCP/oligomycin (FO), and lysed after 2h. pUb levels were measured by ELISA and shown as raw absorbance units. (E) YPMK (WT or PINK1 KO) cells were treated with MTK458 and 1 µM FO, immediately followed by live cell imaging. The percentage of cells with colocalization of YFP-Parkin with mitochondrial mKeima was assessed at each timepoint. (F) YPMK (WT or PINK1 KO) cells were treated with MTK458 (or kinetin) and 1 µM FO for 6h and then analyzed by FACS to measure the percent of cells undergoing mitophagy. (G) YPMK cells were treated with or without MTK458 (3.1 µM) ± 0.5 µM FO for 2h and then collected for mitochondrial isolation. Mitochondrial PINK1 levels and pUb were assessed by immunoblot. (H) SK-OV-3 cells were treated with MTK458 ± FO for 6h and then harvested and analyzed for pUb content by mass spectrometry. (I) Schematic for nanoBRET assay. The nanoBRET assay assesses whether a small molecule (labeled with a BODIPY dye) binds to a particular protein (tagged with NanoLuc). Only when the drug binds to the protein will there be energy transfer between the NanoLuc and the nanoBRET dye, generating a BRET signal. (J) HEK293 PINK1-KO cells were transfected with plasmids encoding PINK1 N-terminally tagged with nanoLuc (NL-PINK1), or GSK-3b C-terminally tagged with NanoLuc (GSK-3b-NL). Cells were then treated with MTK458 labeled with the nanoBRET 590 dye (MTK458 tracer). A high nanoBRET signal was observed with NL-PINK1, but not GSK-3b-NL, suggesting specific binding between PINK1 and MTK458. For all graphs with error bars, mean and SD is graphed. Mean ± SD. *p < 0.05, **p < 0.01, ***p < 0.001, n.s., not significant.

We tested MTK458 in successive assays for PINK1 pathway activity in the presence of low concentrations of FO. In HeLa cells expressing YFP-Parkin and mito-Keima (YPMK), MTK458 increased pUb as assessed by a custom pUb ELISA assay (Figure 3D), Western blotting (Figure 3G), and mass spectrometry (Figure 3H). Next, we monitored Parkin activation by PINK1 though its cytosolic-to-mitochondrial translocation via live-cell imaging. MTK458 accelerated the mitochondrial localization of YFP-Parkin to the mitochondria in a dose-dependent manner (Figures 3E and S4E-F). Thus, MTK458 increases early stage (pUb), mid-stage (Parkin recruitment to mitochondria), and late-stage (mitophagy, Figure 3F) processes of the PINK1/Parkin cascade. In all three assays, we show that MTK458 works in a dose-dependent and PINK1-dependent manner.

The only confirmed substrate of PINK1 is Ubiquitin, but it was reported that PINK1 activation results in phosphorylation of some Rab proteins, specifically Rab8A, 8B and 13, at the highly conserved residue of serine 111 (Lai et al., 2015). The phosphorylation of the Rabs is not catalyzed by PINK1 directly, but is abolished in PINK1 knockout cells, indicating this phosphorylation site can be used as a proxy for PINK1 activity. Consistent with being a PINK1 activator, MTK458 also increases the pS111 Rab8A signal in cells treated with a low dose of FO (Figure S4G-I).

### MTK458 shows direct PINK1 binding

Although PINK1 from non-mammalian species has been utilized for structural studies (Gan et al., 2022), purification of human PINK1 for direct binding assays or crystallization has not yet been achieved. In order to demonstrate direct PINK1 engagement we developed a novel direct binding assay for PINK1 in human cells based on the nanoBRET (bioluminescence resonance energy transfer) system (Machleidt et al., 2015), using a tracer molecule based on the structure of MTK458 (Figures 3I-J). In this system, PINK1 was N-terminally tagged with NanoLuc luciferase (NL-PINK1), and MTK458 was labeled with the nanoBRET 590 dye. With this approach, a BRET signal only results if the labeled MTK458 is within 100 angstroms of NL-PINK1 (Figure 3I). We observed a concentration dependent increase in BRET signal in cells expressing NL-PINK1 and treated with the MTK458-derived nanoBRET tracer (Figure 3J), suggesting that MTK458 binds directly to PINK1. MTK458 did not bind an unrelated protein, GSK3B-NL (Figure 3J). As a control, a non-specific kinase binding tracer K8 (Promega) induced a dose-dependent increase in BRET ratio with the GSK3B-NL, and less signal with NL-PINK1 (Figure S4J), suggesting that MTK-458 promotes mitophagy through direct binding of PINK1.

### MTK458 stabilizes the PINK1/TOM complex and opposes PINK1 inactivation

We next explored the mechanism by which MTK458 potentiates PINK1 activity. Previous work with kinetin and the active metabolite KTP suggested that modification to a triphosphate form is required for activity (Hertz et al., 2013). However, MTK458 cannot be ribosylated (data not shown), so despite similarities in structure, we postulated that MTK458 must act via a new mechanism. Activation of PINK1 involves dimerization, auto-phosphorylation in trans at Ser228, and formation of a high molecular weight (HMW) complex with components of the mitochondrial translocase of the outer membrane (TOM) proteins (Lazarou et al., 2012; Okatsu et al., 2012; Rasool and Trempe, 2018). To investigate the effect of MTK458 on PINK1 dimerization, we used a split-nanoLuc protein fragment complementation system whereby cells were transfected with two species of PINK1, one fused with SmBiT and the other with LgBiT (Figure 4A). When PINK1 dimerizes, the SmBiT and LgBiT proteins assemble into a functional nanoLuc protein that can generate a luminescence signal. MTK458 increased PINK1 dimerization in a concentration dependent manner (Figure 4B). Importantly, MTK458 alone or low-dose F/O alone (t=0 point is +F/O) did not stimulate dimerization, only when both were added was luminescence observed. We used Phos-tag SDS-PAGE (Kinoshita et al., 2009) and blue-native gel electrophoresis (Lazarou et al., 2012) to test the effect of MTK458 on PINK1 phosphorylation and complex formation, respectively. We observed an increase in a phospho-PINK1 species by MTK458 in Phos-tag SDS-PAGE (Figures 4C-E and S5A-C). Also, addition of MTK458 increased the amount of PINK1 in the active, HMW complex (Figure 4F-H).

**Figure 4:**
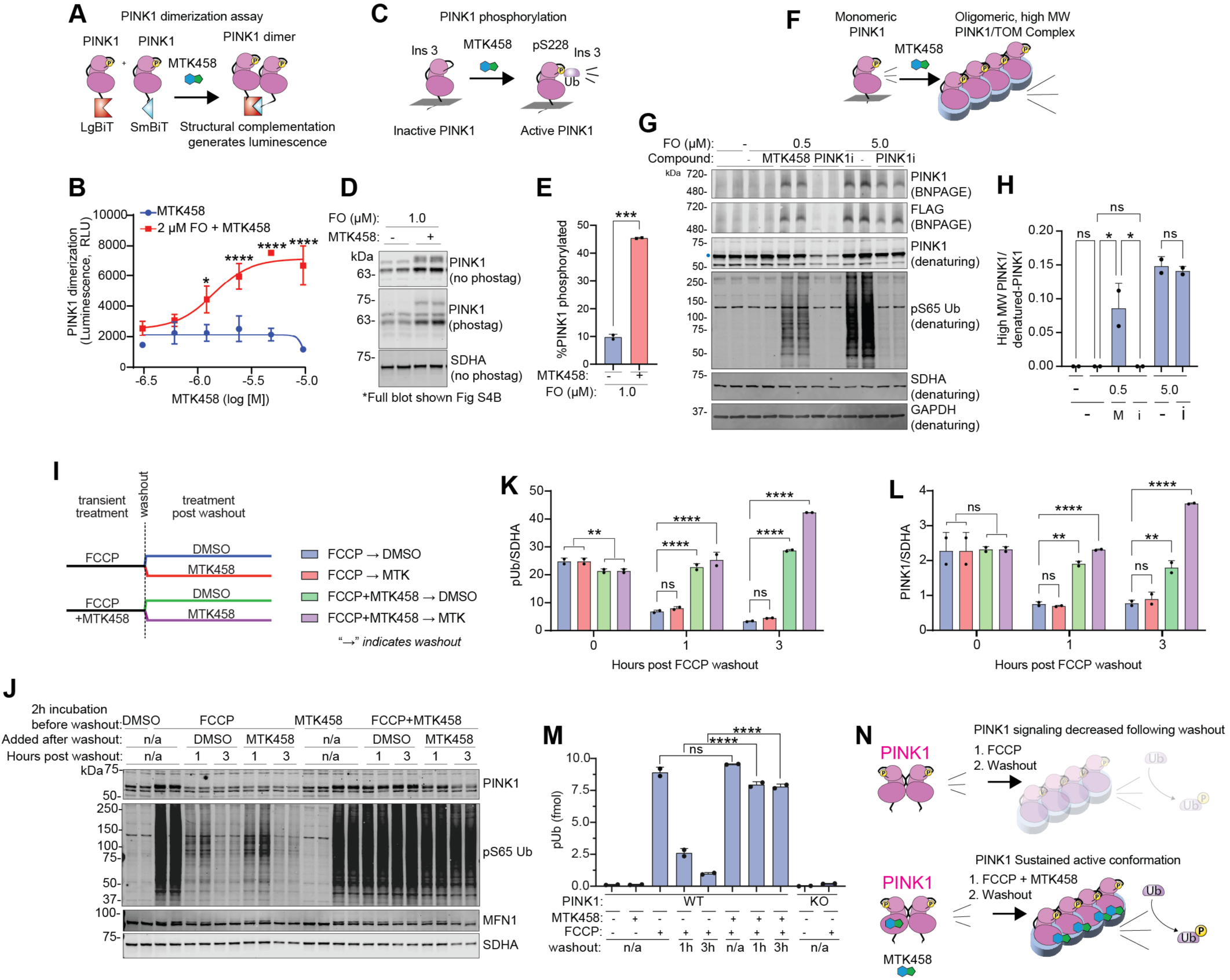
MTK458 stabilizes and sustains the active form of PINK1. (A) Schematic of PINK1 dimerization (nanoBiT) assay. Plasmids encoding PINK1 tagged with either LgBiT or SmBiT were transfected into YPMK PINK1 KO cells. The dimerization of PINK1-LgBiT and PINK1-SmBiT brings the LgBiT and SmBiT together, forming an active luciferase enzyme capable of generating a luminescent signal. (B) YPMK PINK1 KO cells expressing PINK1-LgBiT and PINK1-SmBiT were treated with 2 µM FO and MTK458, and dimerization (as luminescence) was measured and plotted. (C) Schematic of PINK1 autophosphorylation at S228 in the active form of PINK1. (D) In EPF1 cells (cells overexpressing PINK1-FLAG) treated with FO and MTK458, there is an increase in the phospho-PINK1 species. (E) Quantification of (D) is shown. (F) Schematic of the active, high molecular weight PINK1/TOM complex. (G) EPF1 cells were treated with FO, 2.8 µM MTK458, and/or 5 µM BIIB057 (a PINK1 inhibitor, or PINK1i) for 2.5 hours. Whole cell lysates were analyzed by immunoblotting on blue native gels (BNPAGE) or denaturing gels. (H) The band intensities from (G) are quantified; M, MTK458; i, PINK1i. (I) A schematic of the FCCP washout experiment in (J-M). SK-OV-3 cells were treated with 10 µM FCCP or 10 µM FCCP + 2.8 µM MTK458 for 2 hours. Cells were washed 3 times to remove FCCP (“washout”), and then media containing either DMSO or MTK458 was added back to the cells. Cells were harvested for analysis just before the washout, or 1 hour or 3 hours after the washout. (J) Immunoblot analysis of cells collected in the FCCP washout experiment, showing sustained PINK1 stability and activity in cells that experienced co-treatment of FCCP with MTK458. (K-L) Band intensities in (J) were quantified and plotted. (M) pUb levels were quantified from cells collected in the FCCP washout experiment by mass spectrometry. (N) A schematic of PINK1 sustained active conformation with MTK458 treatment is shown. Where applicable, mean and SD are shown and one-way ANOVA was used for statistical analysis. Mean ± SD. *p < 0.05, **p < 0.01, ***p < 0.001, n.s., not significant.

Without mitochondrial stress, PINK1 is rapidly destabilized and degraded (Jin et al., 2010). We found that MTK458 does not activate PINK1 without a mitochondrial stressor, whereas it potentiates both PINK1 autophosphorylation and complex formation with low-dose mitochondrial stress. Based on this finding, we hypothesized that MTK458 may stabilize the active PINK1 complex and slow its inactivation. To investigate the effect of MTK458 on PINK1 complex stability after removal of mitochondrial toxins, we performed FCCP washout studies in SK-OV-3 cells, which express endogenous levels of Parkin and downstream components of the PINK1/parkin pathway (Kakade et al., 2022). SK-OV-3 cells were transiently treated with FCCP alone or FCCP+MTK458 for 2h, then the FCCP was removed by washing the cells three times with FBS-containing medium (“washout”). After the washout, the cells were treated with either DMSO or MTK458 (Figure 4I). Exposure to FCCP induced high PINK1 and pUb levels, which rapidly decreased after washout (Figure 4J-M). However, if the cells were co-treated with MTK458 during the transient FCCP treatment, the high PINK1 and pUb levels were sustained after the washout as detected by both immunoblotting and mass spectrometry (Ordureau et al., 2014) (Figure 4J-M). The high molecular weight PINK1 complex was also sustained by MTK458 even after FCCP is removed (Figure S5D-E). Importantly, MTK458 treatment did not interfere with mitochondrial repolarization, suggesting that the PINK1 complex-potentiating effect is not driven by a compound-driven effect on mitochondrial membrane potential (Figure S5F-G). Taken together, our data supports a model where MTK458 potentiates and prolongs the stability of the active PINK1 complex (Figure 4N), but does not initiate complex stabilization absent mitochondrial depolarization.

### MTK458 drives clearance of pathologic o-synuclein in vitro

We observed mitochondrial dysfunction coupled with increased PINK1 activity in primary neurons (Figure 1K) and so wanted to test whether MTK458 would increase PINK1 activity in models of proeteinopathy. To test this, we utilized two independent in vitro assays, an inducible mitochondrial proteinopathy model and the aggregated α-synuclein seeding models (PFFs) noted above.

First, we employed the inducible cell-based model, which features a deletion mutant of ornithine transcarbamylase (ΔOTC) (Figure 5A) (Moisoi et al., 2014). This mutant yields detergent-insoluble mitochondrial ΔOTC protein aggregates in the mitochondrial matrix that can be cleared by PINK1/Parkin-mediated mitophagy (Burman et al., 2017; Moisoi et al., 2014). In HeLa cells expressing doxycycline-induced ΔOTC and YFP-Parkin, enhancing PINK1 activity with MTK458 treatment resulted in the robust clearance of ΔOTC as measured by either immunofluorescence (Figure 5B-C) or Western blotting (Figure 5D-E), demonstrating that clearance of intramitochondrial aggregates can be enhanced by increased PINK1-mediated mitophagy driven by MTK458.

**Figure 5:**
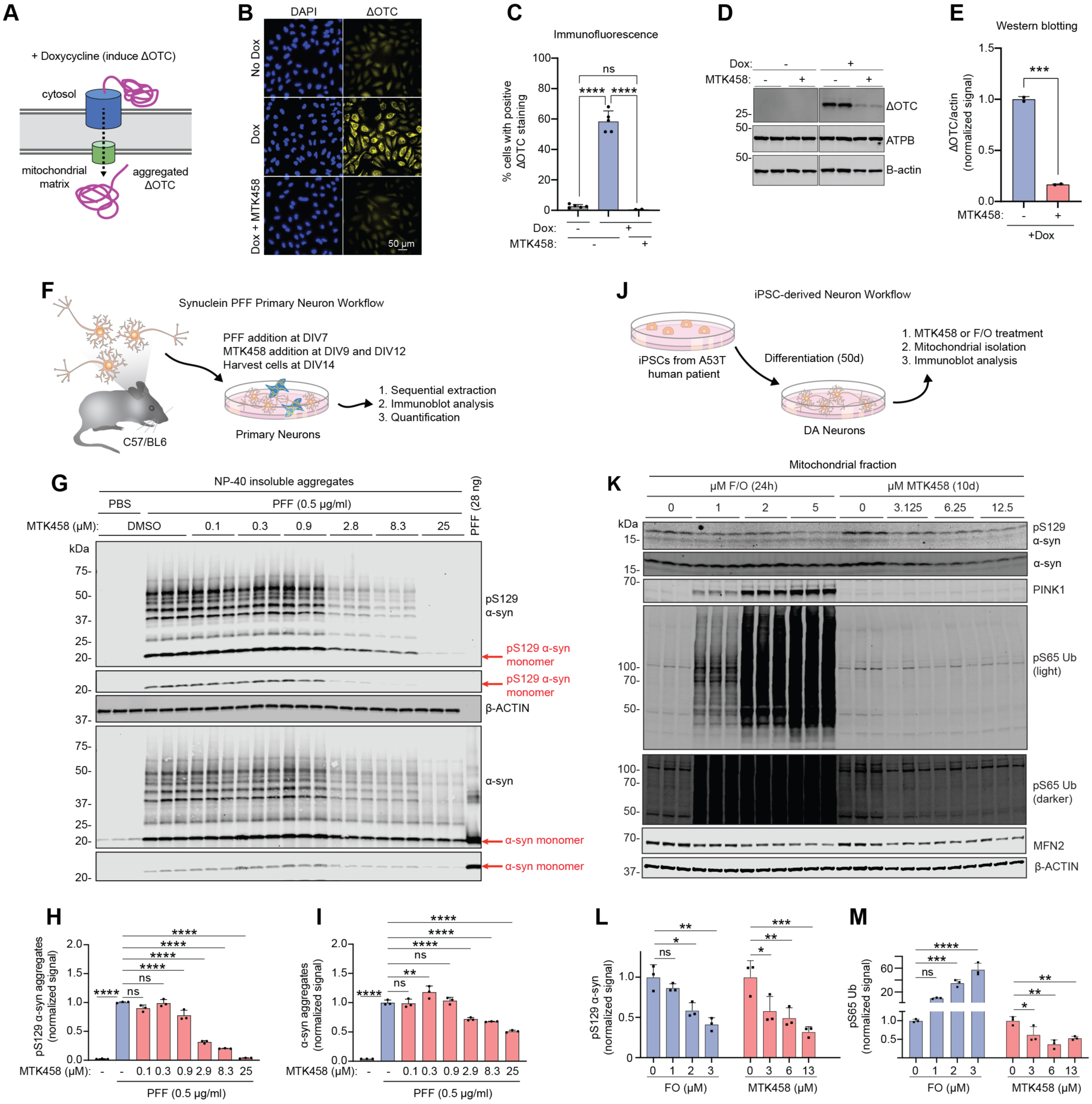
PINK1 activator MTK458 rescues proteotoxicity and PD pathology in immortalized cells, primary neurons, and iPSCderived neurons. (A) Schematic describing the inducible cell-based model expressing a deletion mutant of ornithine transcarbamylase (ΔOTC). This mutant yields detergent-insoluble protein aggregates in the mitochondrial matrix. (B-E) In HeLa cells expressing doxycycline induced ΔOTC and YFP-Parkin, MTK458 treatment (25 µM) results in the clearance of ΔOTC, as assessed by immunofluorescence (B-C) or immunoblotting (D-E). (F) Schematic describing the primary neuron culture experiments in (G-I). Mouse primary cultured neurons were challenged with PFFs (0.5 ug/mL) on DIV7, treated with MTK458 on DIV9 and DIV12, and harvested on DIV14. (G-I) Following serial extraction, the amount of pS129 α-syn in the NP-40 insoluble fraction was quantified from immunoblots, showing a dose dependent decrease in pS129 α-syn aggregates or α-syn aggregates (12-250 kDa) species by MTK458. (J) Schematic describing the iPSC-derived neuron workflow (K) Human A53T patient-derived iPSC neurons were treated for 10 days with MTK458 or 24h with FO, and harvested and analyzed by immunoblotting. (L-M) Quantification of (K) is shown. Mean ± SD. *p < 0.05, **p < 0.01, ***p < 0.001, n.s., not significant.

Having shown that MTK458 treatment could reduce mitochondrial aggregates in an inducible model in immortalized cells, we next tested whether PINK1 activation could ameliorate PD-patient relevant pathology in mouse and human neurons in vitro. Primary mouse hippocampal neuron cultures seeded with α-synuclein PFFs on DIV7 (Volpicelli-Daley et al., 2014) were allowed to seed and further develop detergent insoluble pS129 α-synuclein aggregates prior to the addition of MTK458 on DIV9 and DIV12 (Figure 5F). On DIV14, pS129 α-synuclein aggregates were detectable by immunoblotting from the insoluble fraction (Figure 5G) and immunofluorescence (Figure S6A). MTK458 treatment led to the clearance of pS129 α-synuclein aggregates (12-250 kDa) in a dose-dependent manner (Figures 5G-I).

We further tested the effect of MTK458 in iPSC-derived neurons from patients with the A53T-α-synuclein mutation associated with PD (Figure 5J). This line carries an A53T α-synuclein mutation that causes the derived DA neurons to accumulate pS129 α-synuclein without the addition of exogenous PFFs, and additionally serves to bridge primary mouse neuron studies with human neurons. To first test if global induction of mitophagy driven could reduce pS129 α-synuclein, cells were treated with FCCP to activate PINK1 by depolarization (Figures 5K-M). FCCP alone reduced pS129 α-synuclein pathology in these cells, but at the expense of an increase in mitochondrial stress throughout the cell, as evidenced by stabilization of PINK1 and increased pUb levels (Figures 5K-M). In contrast, MTK458 treatment for 10 days reduced α-synuclein pathology and the mitochondrial stress marker pUb (Figures 5K-M). We hypothesize that PINK1 stabilization does happen in cells treated with MTK458 alone, but at lower levels and selectively on impaired mitochondria. Consistent with the model we proposed above, activation of PINK1 in the patient-derived iPSC neurons reduced protein aggregate load and ultimately drove a reduction in pUb (Figures 5K-M). Our results in various PD cell models suggest that pharmacological augmentation of PINK1 activity can ameliorate proteinopathy.

### MTK458 drives clearance of pathologic α-synuclein in vivo

We next tested whether MTK458 could rescue α-synuclein pathology in mice. We utilized a widely adopted pre-clinical model for PD in which α-synuclein PFFs are injected unilaterally into the striatum of mice, leading to progressive, spreading α-synuclein pathology (Figure 6A). MTK458 has excellent mouse pharmacokinetics and high brain penetrance. A microdialysis study in the mouse striatum showed a similar unbound plasma and brain exposures at equilibrium (unbound partition coefficient, Kp _u,u_ ∼1) (data not shown). Consistent with our finding in mouse primary neuron cultures that PFF incubation led to an increase in α-synuclein pathology and concurrent increase in pUb, we found that PFF injection into the striatum robustly induces aggregated and pS129 α-synuclein after several weeks incubation and an increase in pUb (Figures 6B, 6G, and S6B-D) (Luk et al., 2012). PFF, but not PBS injection, increases brain pUb (Figure 6G), and central (TREM2) and peripheral (IL6, CXCL1) inflammatory markers (Figure S6E-G). Daily oral administration of MTK458 in these mice led to dose dependent decrease (up to ∼50%) in α-synuclein pathology in 3-month, and 7-month studies (Figures 6B-D and S6H-J). MTK458 also rescued an activity deficit in freely moving PFF mice as assessed by home cage monitoring (Figures 6E-F). The dose-response rescue in pathology matched the rescue in motor activity. The increase in TREM2, IL-6, and CXCL1 inflammatory markers were also attenuated by MTK458 (Figures S6E-G), which is in line with a role for PINK1 in inflammation (Sliter et al., 2018).

**Figure 6:**
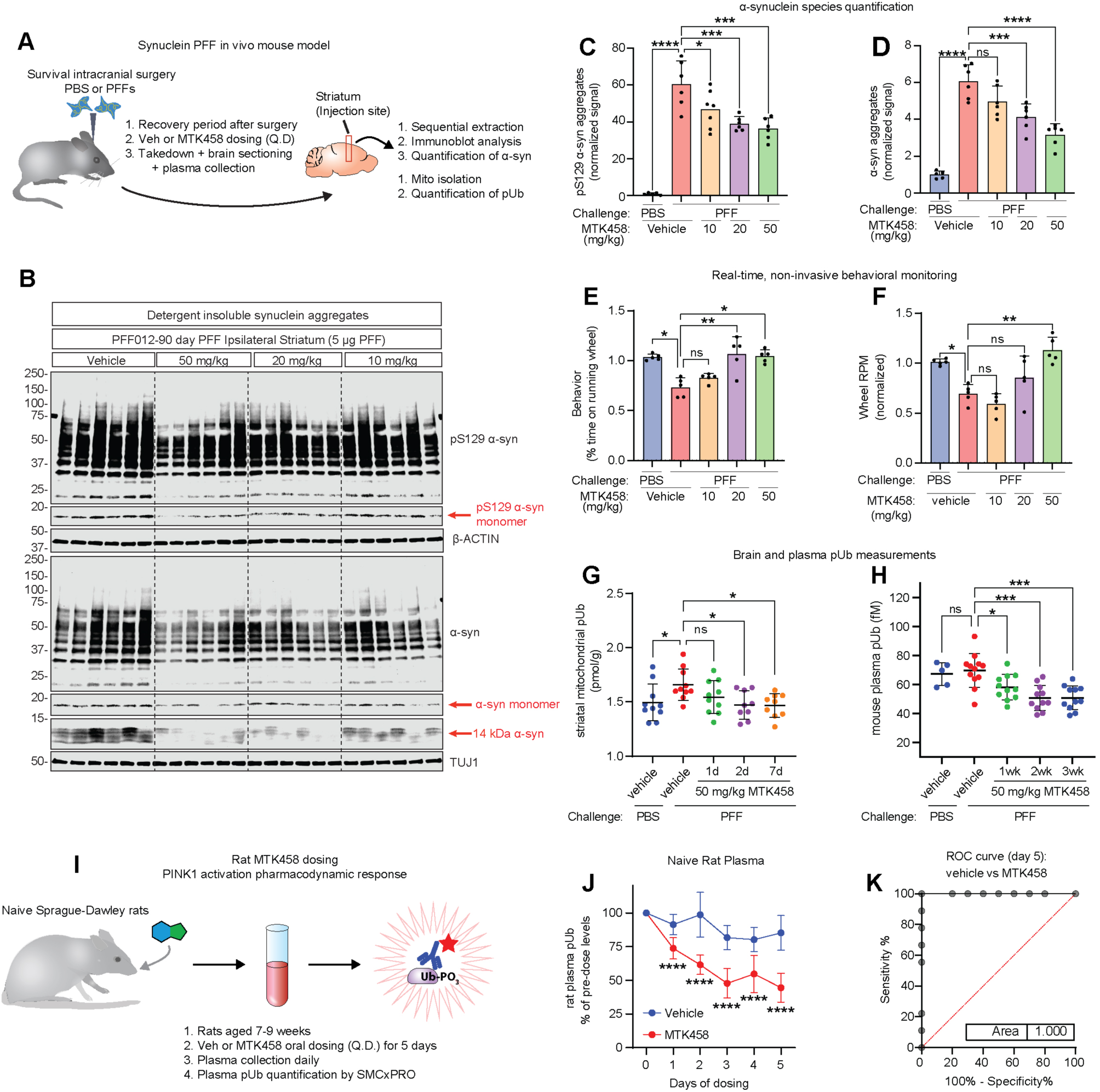
PINK1 activator MTK458 rescues PD pathology and normalizes p-S65-Ub levels in vivo. (A) Schematic of *in vivo* PFF experiments in mice. Mice were challenged with striatal injection of PFFs and dosed (QD, PO) with MTK458 at the indicated concentrations. (B) After 90 days, striatum brain pieces were analyzed for aggregate pS129 α-syn or total α-syn species (12-250 kDa). (C-D) Quantification of analysis of the full cohort is shown here. (E-F) Parallel mice were challenged with striatal injection of PBS or PFF, and then dosed with MTK458 for 6 months. Mice were analyzed for time spent on the running wheel using the Vium system. (G) Mice were challenged with striatal injection of PFFs and after 3 months, were dosed (QD, PO) with 50 mg/kg MTK458 for the indicated times. Mitochondria was isolated from striatal brain pieces and analyzed for pUb content on the MSD assay. (H) Mice were challenged with PFFs, and after 3 months, were dosed (PO, QD) with 50 mg/kg MTK458 for the indicated durations. Plasma pUb levels were measured by the SMCxPRO pUb assay. (I) Schematic of rat dosing study to evaluate plasma pUb as a target engagement biomarker for PINK1 activator compounds. (J) Naïve Sprague-Dawley rats were dosed (PO, QD) with 50 mg/kg MTK458 for 5 days (6 doses). Plasma pUb levels at the indicated timepoints were determined by the SMCxPRO pUb assay. (K) ROC curve for plasma pUb levels in vehicle or MTK458 dosed rats (after 5 days of dosing) is shown. Mean ± SD. *p < 0.05, **p < 0.01, ***p < 0.001, n.s., not significant.

Consistent with the results from our iPSC-derived neuron experiments, MTK458 treatment decreased pUb in the brains of PFF seeded mice (Figure 6G), suggesting a reduction in mitochondrial stress level due to the clearance of damaged mitochondria and pS129 α-synuclein aggregates. Remarkably, a decrease in plasma pUb was also observed in mice treated with MTK458 (Figure 6H), suggesting that pUb changes in a peripheral biofluid are reflective of brain pUb and pathology. Unexpectedly, we did not see an increase in plasma pUb levels in the PFF-challenged mice as compared to PBS challenged mice.

Since plasma concentrations of pUb in naïve, wild-type animals are measurable and greater than in PINK1 KO animals (Figure S1B-D), we hypothesized that we might be able to detect a decrease in pUb after a short-term MTK458 treatment in animals. Such a change would be useful to measure target engagement of PINK1 activator compounds in animals or in the clinic. In order to test this hypothesis, we dosed naïve, wild-type Sprague-Dawley rats for five days with either a vehicle control or 50 mg/kg MTK458, and found a significant decrease in pUb compared to the vehicle-treated or pre-dosed rats (Figures 6I-J and 7A). The magnitude of the plasma pUb lowering effect (ROC = 1.00) (Figures 6K and 7B-E) suggests it may be useful as a specific and sensitive pharmacodynamic biomarker for MTK458 treatment.

**Figure 7:**
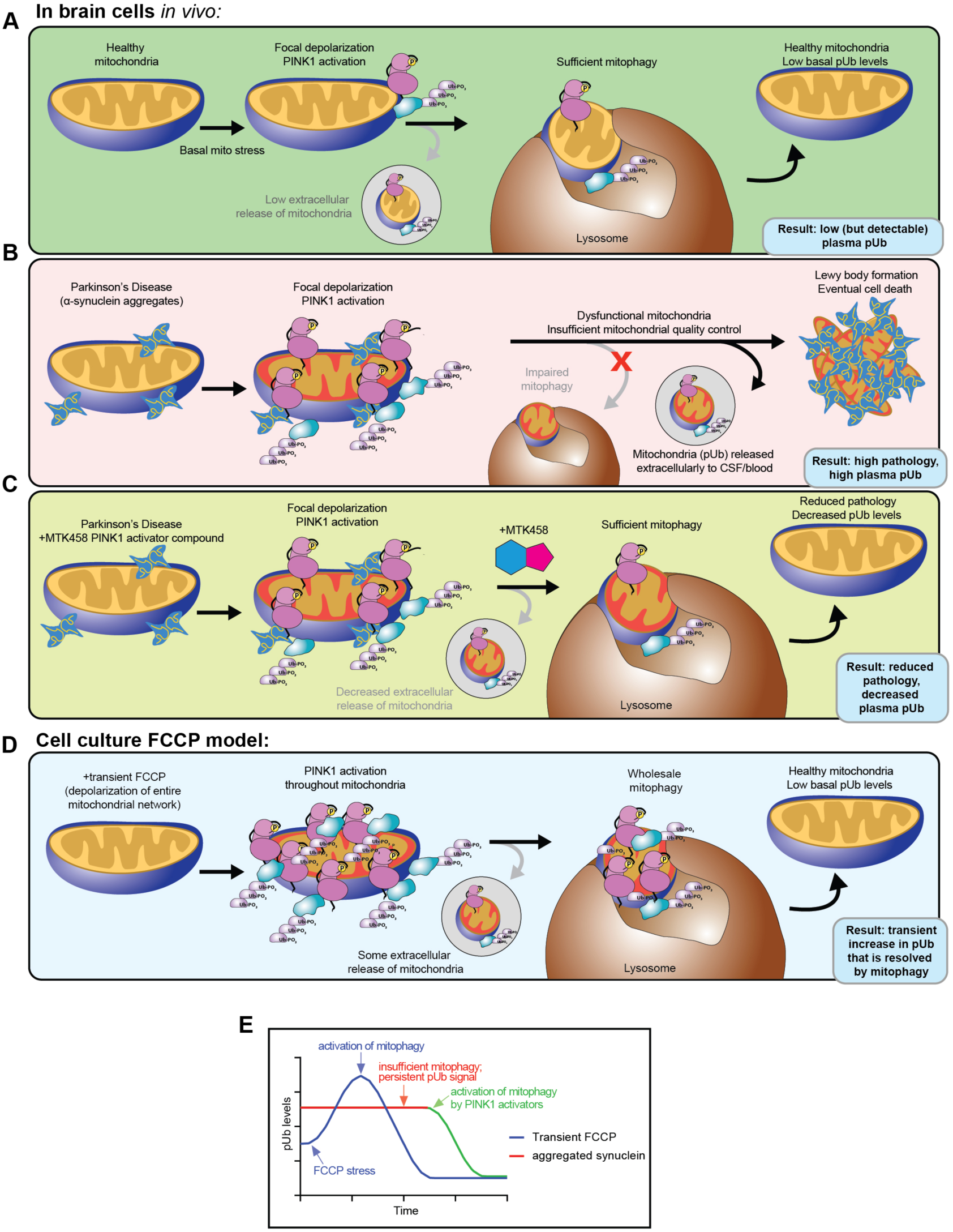
Schematic of pUb regulation. Model for regulation of extracellular pUb levels by mitophagy and therapeutic hypothesis for PINK1 activation. (A) In otherwise healthy mitochondria, stochastic levels of heterogeneous stress can cause focal depolarization of the mitochondrial network, resulting in focal activation of PINK1 and mitophagy of the damaged mitochondria. Basal levels of pUb in the blood are low because few mitochondria are released extracellularly. (B) In PD models, α-synuclein aggregation occurs at the mitochondria, causing focal PINK1 activation and an increase in pUb signal. However, because α-synuclein aggregation inhibits mitophagy, there is an increase in the release of extracellular mitochondria and elevated levels of pUb detected in the brain and the periphery. (C) In PD models treated with MTK458, the potentiated increase in PINK1 activation rescues the mitophagy deficit, leading to reduced pathology and pUb levels. (D) FCCP treatment in cells results in homogeneous mitochondrial depolarization which drives large-scale PINK1 stabilization and pUb upregulation. Eventually, mitophagy will reduce the pUb levels back to normal (or below normal). (E) A schematic of the timecourse of pUb in cells treated with a transient FCCP versus challenged with α-synuclein aggregation. Cells treated with FCCP exhibit a transient increase in pUb that is resolved via mitophagy. Cells challenged with α-synuclein aggregation have sustained high levels of pUb until PINK1 activator drugs are introduced, which activate mitophagy and decrease pUb levels.

## Discussion

In this study, we showed that α-synuclein pathology localizes to mitochondria, causing mitochondrial dysfunction and stalled mitophagy. We also showed that PD patients display higher plasma levels of the PINK1 substrate pUb, which suggests that stalled mitophagy and mitochondrial dysfunction are features of idiopathic Parkinsonism, linking the more common form with rare familial PD associated with mutations in *PINK1.* To address these deficits, we synthesized hundreds of drug-like kinetin analogs and tested their ability to activate PINK1. In this process, we discovered the small molecule MTK458, which binds to and stabilizes the active form of PINK1, increasing its activity and activating mitophagy downstream. In both cellular and animal PD models, PINK1 activation with MTK458 alleviated the hallmark α-synuclein aggregation, mitochondrial dysfunction, and stalled mitophagy that occurs in PD. PINK1 activation also reduced the PINK1-specific biomarker pUb, an indicator of mitochondrial stress that correlates with the severity of PD in humans. Our data serve as a preclinical proof-of-concept supporting PINK1 activation as a strategy for treating PD.

### PINK1 as a target for disease modification

PINK1 has several features that make it inherently attractive as a drug target. First, LoF mutations in PINK1 are genetically linked to Parkinson’s disease. Second, mitochondrial dysfunction is associated with PD and PINK1 activity plays a central role in triggering stress-related mitochondrial quality control processes, suggesting that PINK1 activation could have a disease-modifying benefit (Borsche et al., 2021; Kim et al., 2021). Third, PINK1 has an endogenous regulatory mechanism that limits its presence only to conditions involving mitochondrial stress, so pharmacological activation of PINK1 should not constitutively activate kinase activity. An alternative target could be the E3-Ubiquitin ligase Parkin, which is also genetically linked to PD. However, Parkin is a less attractive therapeutic target because it is present constitutively, can affect proteasome-mediated degradation, and is difficult to activate pharmacologically (Shlevkov et al., 2022). Our group and others previously published that the neo-substrate kinetin tri-phosphate (KTP) could be used as an alternative phospho-donor by PINK1 with higher catalytic efficiency than ATP, and that the pro-drug kinetin could be taken up by cells and converted to KTP (Hertz et al., 2013; Osgerby et al., 2017). Others have shown that kinetin can activate PINK1 and rescue the mitochondrial mutation load and climbing activity in heteroplasmic flies in a PINK1-dependent manner (Tsai et al., 2022). However, kinetin is not developable due to low potency, low brain penetration, and unfavorable pharmacokinetics, which limited its efficacy in mammalian in vivo models (Orr et al., 2017).

Using kinetin as a starting point, we searched for more potent molecules with attractive drug-like properties following a structure-activity-relationship (SAR)-driven approach. We screened compounds using both mitophagy (efficacy) and mitochondrial toxicity (safety) assays to eliminate potential hits that were simply mitochondrial toxins. We then confirmed that the active compounds resulted in the activation of the entire PINK1 signaling cascade, including Ub phosphorylation, Parkin recruitment, and mitophagy. We identified MTK458 as a PINK1 activator that directly binds to PINK1 and stabilizes the active form of PINK1. MTK458 retains the mitophagy activating properties of kinetin but has enhanced pharmacokinetics, including brain penetration and improved metabolic and physiochemical properties. Using MTK458 to activate mitophagy restored cellular quality control capabilities for mitochondrial homeostasis and resulted in reduced levels of pathogenic α-synuclein *in vitro* and *in vivo.* Despite being observed by several labs (Butler et al., 2012; Lee et al., 2002; Nakamura et al., 2011; Ryan et al., 2018; Shaltouki et al., 2018), the detailed mechanism of how mitophagy and improved mitochondrial homeostasis lead to mitochondrial protein aggregate clearance is not known. The amelioration of α-synuclein pathology may be a direct result of clearing the mitochondria associated α-synuclein or an indirect result of the increased mitochondrial homeostasis, which would stabilize ATP levels leading to more lysosomal and proteosomal activity for the overall improvement in cellular proteostasis.

### pUb as a biomarker for PD

Human genetics established the *PINK1/PRKN* pathway as being causative in PD (Kitada et al., 1998; Valente et al., 2004). However, the pathogenic mechanisms by which LoF mutations lead to disease are not well understood, and the role of the *PINK1/PRKN* pathway in idiopathic disease is unknown. PINK1 is the only known Ubiquitin kinase, so pUb serves both as a specific biomarker of PINK1 activity and a general biomarker for mitochondrial health. We discovered that plasma pUb levels are elevated in PD patients and correlate with clinical progression scores, positioning it as a candidate disease progression biomarker. As a diagnostic biomarker, pUb has similar predictive value to elevated LDL cholesterol for cardiac mortality (ROC=0.67 pUb in PD, ROC=0.63 LDL cholesterol for cardiac mortality (Doran et al., 2014). The elevated plasma pUb levels in idiopathic PD patients confirm that mitochondrial dysfunction is a general feature of PD, rather than occurring only in PD cases associated with mitochondrial mutations. Based on these data, future work should focus on analyzing longitudinal samples from individual patients to determine whether plasma pUb could serve as a disease progression biomarker in clinical trials.

### PINK1 substrate pUb biomarker mechanism

Our finding that pUb is elevated in PD patients and that a PINK1 activator reduces pUb may seem unexpected. However, it is consistent with several features of our selective, PINK1-targeted strategy and the interplay between α-synuclein pathology and mitochondrial dysfunction. Our model is shown in Figure 7. In healthy mitochondria, basal mitochondrial stress causes focal, localized depolarization of the mitochondria (Figure 7A). PINK1 is stabilized in these depolarized regions of mitochondria, resulting in fragmentation, engulfment of the damaged piece of mitochondrial network by autophagic machinery, and restoration of mitochondrial health. When mitophagy is not compromised, only low, basal amounts of mitochondria are released extracellularly, which results in relatively low levels of pUb detected in the plasma in healthy patients/animals. In PD, α-synuclein aggregates localize to the mitochondria, causing focal depolarization, PINK1 activation, and an increase in pUb (Figure 7B). Normally, pUb spikes would be quickly resolved by mitophagy. However, because mitophagy is compromised by α-synuclein aggregation (Figure 2J) (Shaltouki et al., 2018), the increase in pUb is stabilized and detectable in cells and tissues (Figures 2K and 6G). Mitophagy impairment leads to an increase in the release of mitochondrion (containing pUb) extracellularly (Choong et al., 2021; Tan et al., 2022), which results in elevated levels of plasma pUb. Ultimately, if the pathology remains unresolved, α-synuclein aggregation leads to Lewy body formation and eventual neuronal death. This pathology can be rescued with compounds that activate PINK1, such as MTK458. As we showed in preclinical models of PD, treatment with a PINK1 activator rescued mitophagy, decreased α-synuclein aggregation, and lowered pUb (Figures 7C and 7E).

In comparison, mitochondrial toxins like FCCP causes depolarization of the entire mitochondrial network, leading to a more global activation of PINK1 throughout the network and a significant increase in pUb. Mitophagy degrades most of the depolarized mitochondria, with some being released extracellularly, and only once the toxin is removed is the transient pUb spike is resolved (Figures 7D-E).

## Summary

In conclusion, our data support the idea that increasing PINK1 activity to induce mitophagy could be a viable therapeutic approach for disease-modification in idiopathic PD. Data from our group and others suggest a model whereby α-synuclein pathology causes mitochondrial dysfunction and suppresses mitophagy, thereby increasing pUb. The novel small molecule MTK458 binds to PINK1 and stabilizes its active complex, triggering the first step in mitophagy. In both cellular and animal models of α-synuclein aggregation (PD-like pathology), MTK458 decreased pS129 α-synuclein aggregates and normalized both brain and corresponding plasma pUb levels. PINK1 activation may thus be sufficient to address the hallmark α-synuclein pathology observed in PD and the resultant mitochondrial dysfunction. Altogether, our data demonstrate that PINK1 activators can rescue pathology associated with idiopathic PD and that this class of molecules is worthy of further preclinical and clinical exploration as potential disease modifying therapeutics for PD.

## Acknowledgments

We would like to thank the MJFF team, in particular Dr. Shalini Padmanabhan and Dr. Samantha Hutten, for providing human plasma samples from PD or HC patients for pUb analysis. Data used in preparation of this article were obtained from the MJFF-sponsored LRRK2 Cohort Consortium (LCC). For up-to-date information on the study, visit www.michaeljfox.org/lcc. The LRRK2 Cohort Consortium is coordinated and funded by The Michael J. Fox Foundation for Parkinson’s Research. PINK1 and Parkin double knockout rat samples were provided by Dr. Kelly Stauch (University of Nebraska). Plasma samples from humans containing PINK1 or Parkin mutations were provided by Dr. Christine Klein (University of Lubeck). YPMK and ΔOTC cell lines were provided by Dr. Richard Youle (NIH). Plasmids containing PINK1 used in the nanoBRET and nanoBIT assays were provided by Promega (Steven Edenson, Matt Robers). We also acknowledge Henrietta Lacks and her family for providing the HeLa cell line used in our work. This work was supported in part by Michael J. Fox Foundation grant MJFF-001839 to N.T.H., Michael J. Fox Foundation grant 15866 to J.W.H and NIH grant NS083524 to J.W.H.

## Author Contributions

Conceptualization, R.M.C., R.R., D.D., and N.T.H.; Methodology, R.M.C., R.R., D.D., C.W., and N.T.H; Investigation, R.M.C., R.R., D.D., C.W., S.L., A.M., A.L., A.E., C.T., R.Y.K., J.L., S.H., S.K., V.R., D.H., V.G., J.P., A.O., and N.T.H.; Writing – Original Draft, R.M.C., R.R., D.D., and N.T.H.; Writing – Review & Editing, R.M.C., R.R., D.D., J.S., Crystal Herron (Redwood Ink), and N.T.H.; Visualization, R.M.C., R.R., D.D., and N.T.H.; Supervision, R.M.C., R.R., D.D., W.H., S.J., J.S., and N.T.H.; Project Administration, D.d.R. and N.T.H.; Funding Acquisition, D.d.R. and N.T.H.

## Data and Materials Availability

All data needed to evaluate the conclusions of the paper are present in the paper and/or the supplementary materials.

---

**Supplementary Figure S1:**
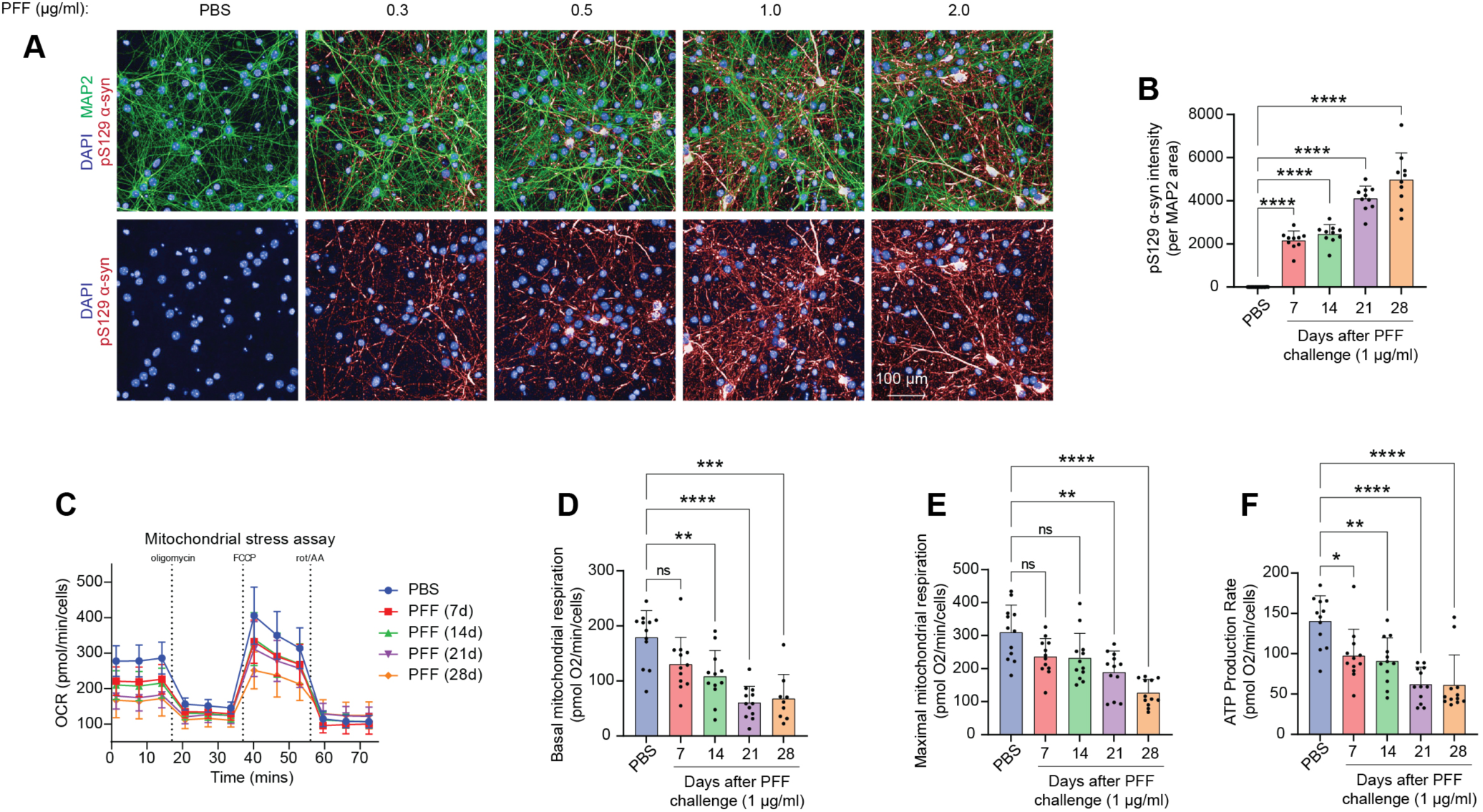
Effects of PFF challenge on mitochondrial respiration and mitophagy in primary neurons. (A-F) M83 primary cultured neurons were challenged with 1 µg/ml of PFFs for the indicated number of days, and pS129 α-synuclein signal (A-B) was assessed by immunofluorescence. (C) Mitochondrial oxygen consumption rate in cell mito stress assay was assessed by the seahorse bioanalyzer, yielding measurements for mitochondrial basal respiration (D), maximal respiration (E), and ATP production rate (F). Mean and SD is shown; one-way ANOVA was used for statistical analysis. Mean ± SD. *p < 0.05, **p < 0.01, ***p < 0.001, n.s., not significant.

**Supplementary Figure S2:**
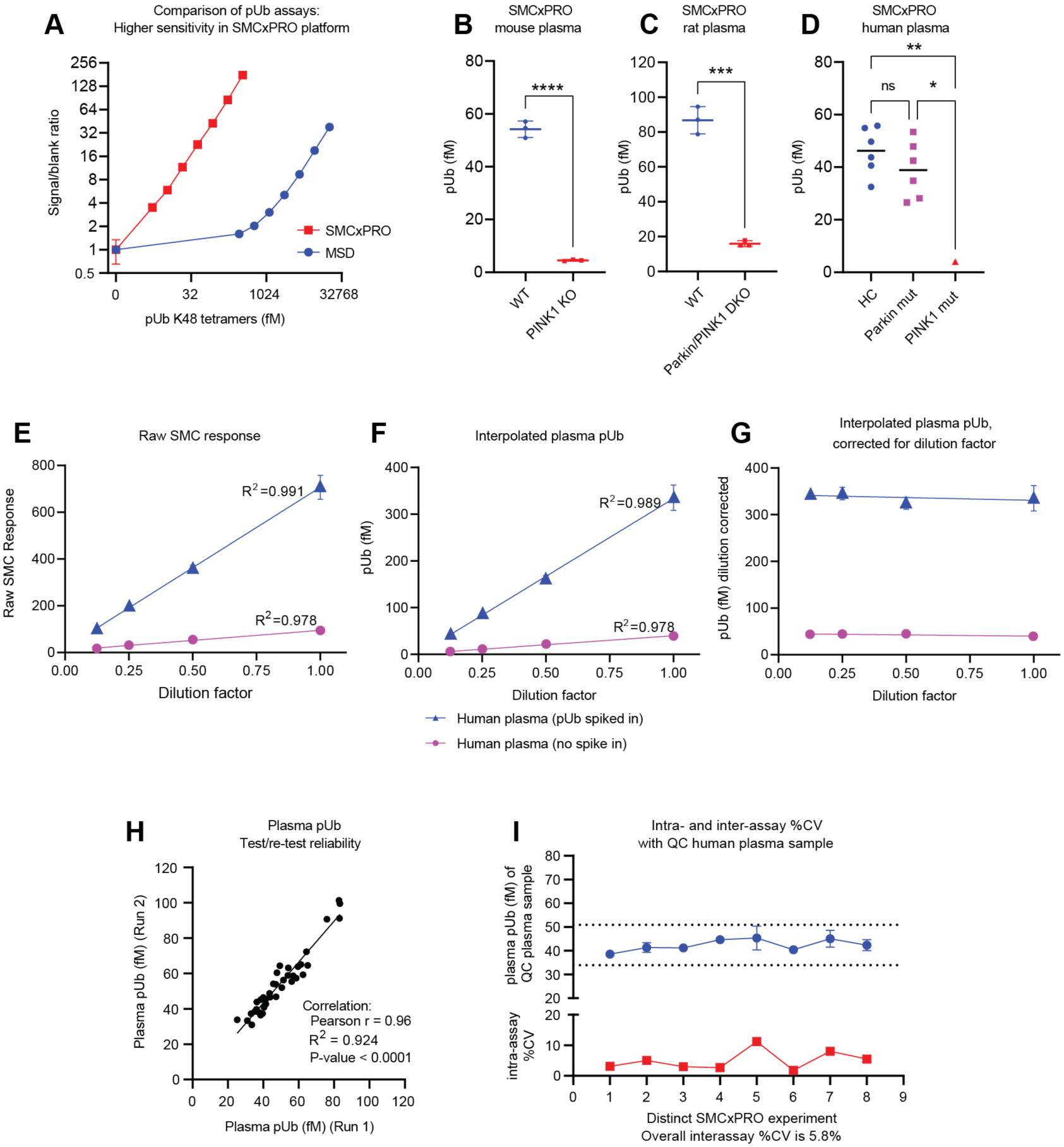
SMCxPRO pUb assay development. (A) Comparison of signal to blank ratio of K48 pUb tetramers on the MSD or SMCxPRO pUb assays. (B-D) If analyzed on the SMCxPRO pUb assay, plasma pUb levels in PINK1 KO mice, rats, or humans are significantly lower compared to WT. (E-G) Assay validation experiments for human plasma on the SMCxPRO pUb assay to assess parallelism, dilution linearity, and spike-recovery (Piccoli and Sauer, 2019). (E) A human plasma sample, spiked in with vehicle or 277 fM pUb, was serially diluted and analyzed on the SMCxPRO pUb assay. The raw SMC responses for the plasma samples at different dilutions are graphed. (F) The raw values in (E) were interpolated to fM of pUb using a calibrator curve of K48 pUb tetramers. (G) Plasma pUb concentrations calculated at each dilution were multiplied by their respective dilution factor and graphed. The relatively flat line confirms that there is good parallelism and dilution linearity in human plasma. The signal in the spiked sample is approximately 277 fM higher than the signal in the non-spiked sample, as expected. (H) A comparison of plasma pUb levels from two independent experiments done on separate days show an excellent test/retest reliability. (I) For a QC human plasma sample ran on multiple plates over multiple days, the interassay CV was below 10% and intraassay CV was below 20%. Unless otherwise indicated, mean and SD is shown; either t-test (for comparison of two groups) or one-way ANOVA (for comparison for 3 or more groups) was used for statistical analysis. Mean ± SD. *p < 0.05, **p < 0.01, ***p < 0.001, ****p<0.0001, n.s., not significant.

**Supplementary Figure S3:**
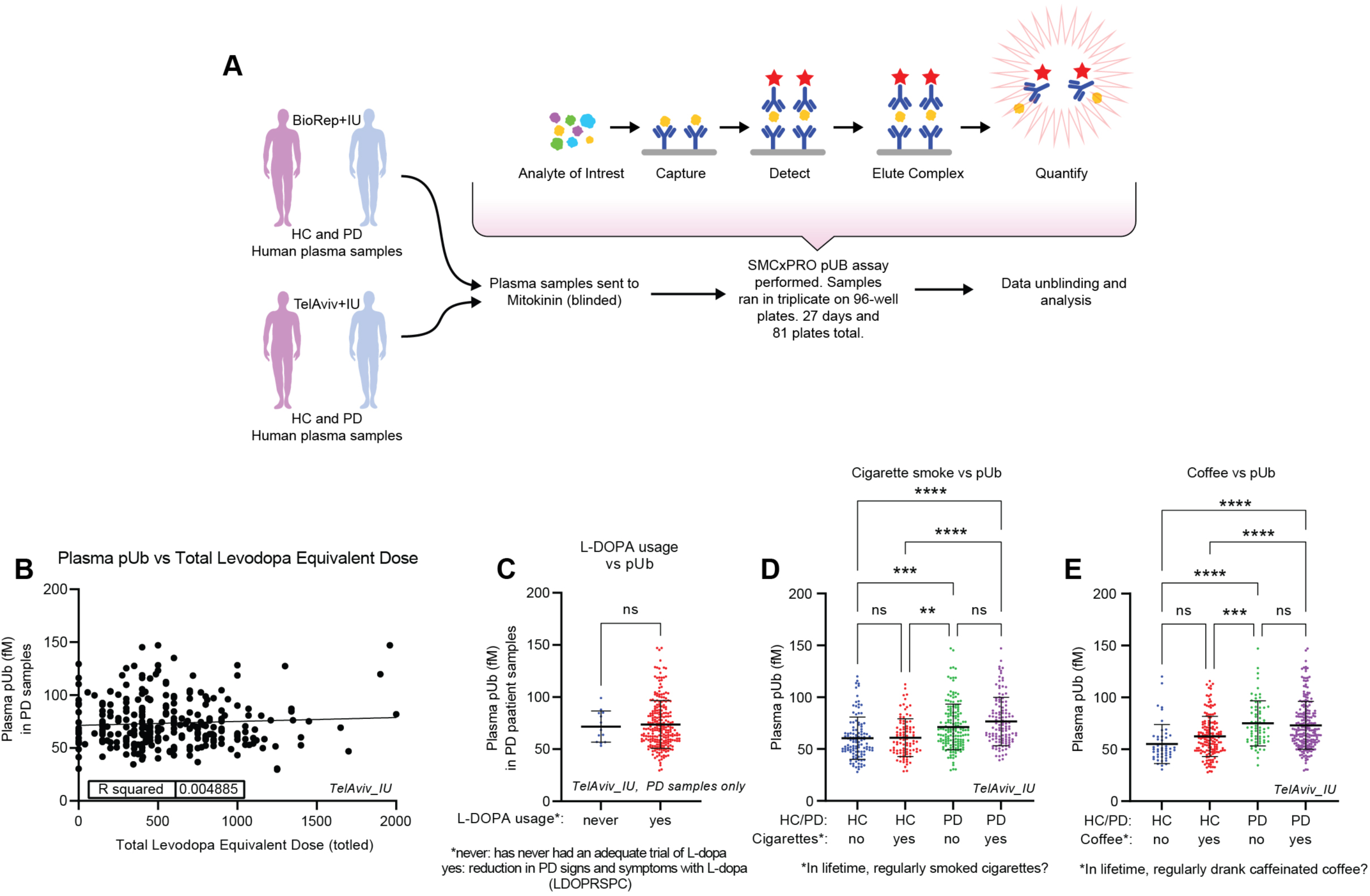
Quantification of pUb levels from in human plasma. (A) Schematic of the analysis of plasma pUb from human HC and PD patients. Human plasma samples were analyzed by the SMCxPRO pUb assay in a blinded fashion. (B) The total levodopa equivalent daily dose (totled) and plasma pUb levels in PD patients are graphed, showing a lack of correlation. (C) Patient plasma pUb levels are binned according to whether they have used L-DOPA or not. There is no significant difference in plasma pUb due to L-DOPA usage. Mean and SD is shown; t-test was used for statistical analysis. (D-E) Patient plasma pUb levels are binned according to whether the patient regularly smoked cigarettes or drank caffeinated coffee. Neither activity significantly changed plasma pUb levels.

**Supplementary Figure S4:**
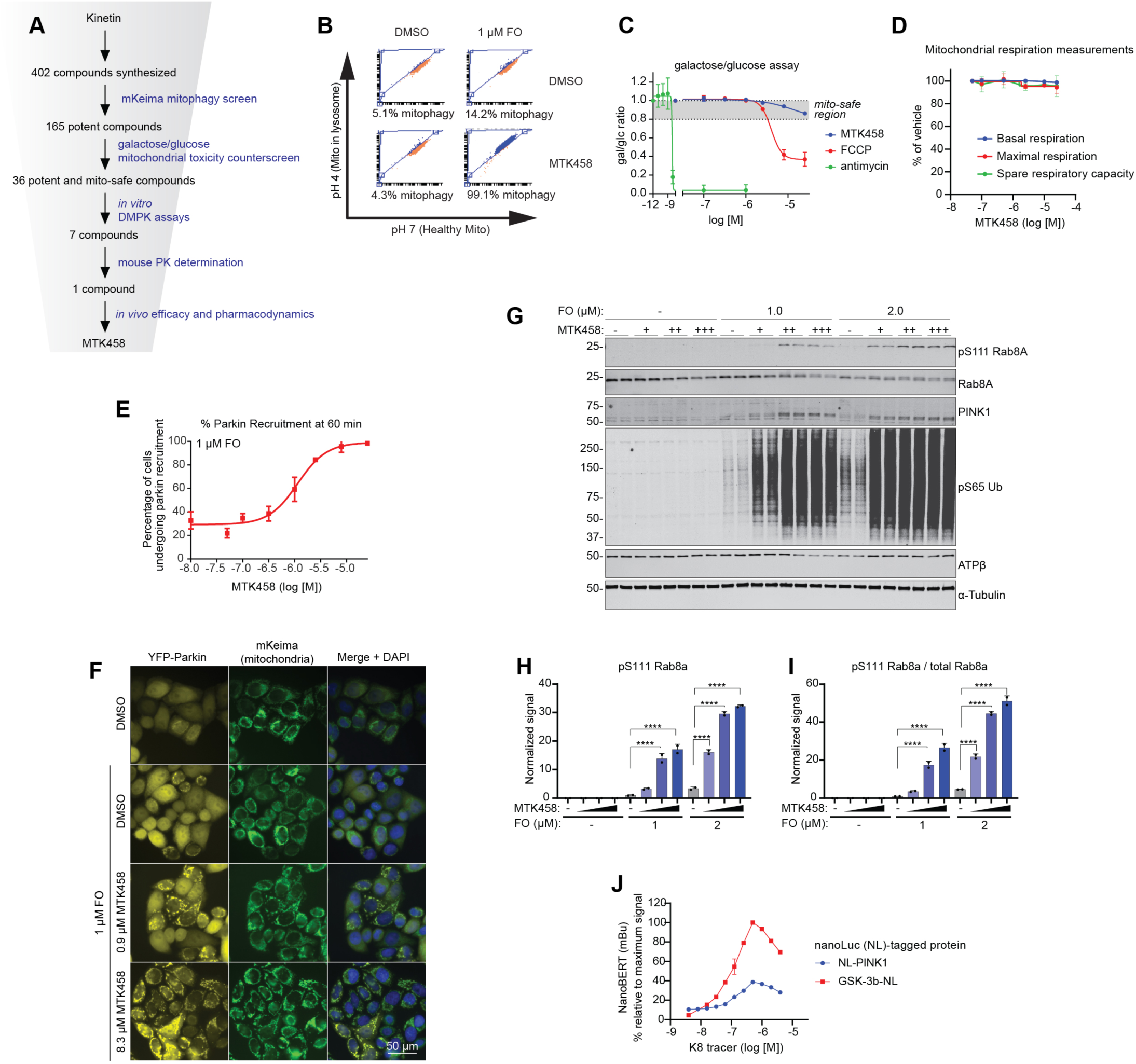
Screening for PINK1 activator compounds. (A) Schematic of the screening funnel used to discover MTK458. (B) Example FACS plots from the mKeima mitophagy screening assay. MTK458 causes an increase in cells undergoing mitophagy if treated in the presence of FO. (C) An example run of the galactose/glucose assay. SK-OV-3 cells grown in galactose or glucose media were treated with MTK458, FCCP, or antimycin. After 24 hours, the ratio of cells remaining in galactose or glucose media (gal/glc ratio) is plotted. An cut off of a 20% decrease in the gal/glc ratio was designated as the threshold for a mitotoxic compound. FCCP and antimycin are mitotoxic, but MTK458 is “mito-safe”. (D) HeLa cells were treated with MTK458 for 1 hour and mitochondrial respiratory measurements were assessed by the Seahorse Bioanalyzer. MTK458 does not cause any impairment of mitochondrial function. (E) YPMK cells were treated with MTK458 and 1 µM FO, immediately followed by live cell imaging. The percentage of cells with colocalization of YFP-Parkin with mitochondrial mKeima was assessed at 1 hour and plotted. (F) Images of YPMK cells in the Parkin recruitment assay at one hour after FO ± MTK458 treatment. The percentage of cells with YFP-Parkin localized to the mitochondria is increased with MTK458 treatment. Scale bar, 50 µm (G) SK-OV-3 cells were treated with increasing doses of MTK458 in conjunction with a low dose of FO. MTK458 treatment dose-dependently increases the PINK1 dependent pS111-Rab8A signal, as shown by Western blotting. (H-I) Quantification of (G). (J) As a negative control for Figure 3J, we used a proprietary (Promega, K8) tracer that binds to GSK-3b-NL, but not PINK1. Mean ± SD. *p < 0.05, **p < 0.01, ***p < 0.001, ****p<0.0001, n.s., not significant.

**Supplementary Figure S5:**
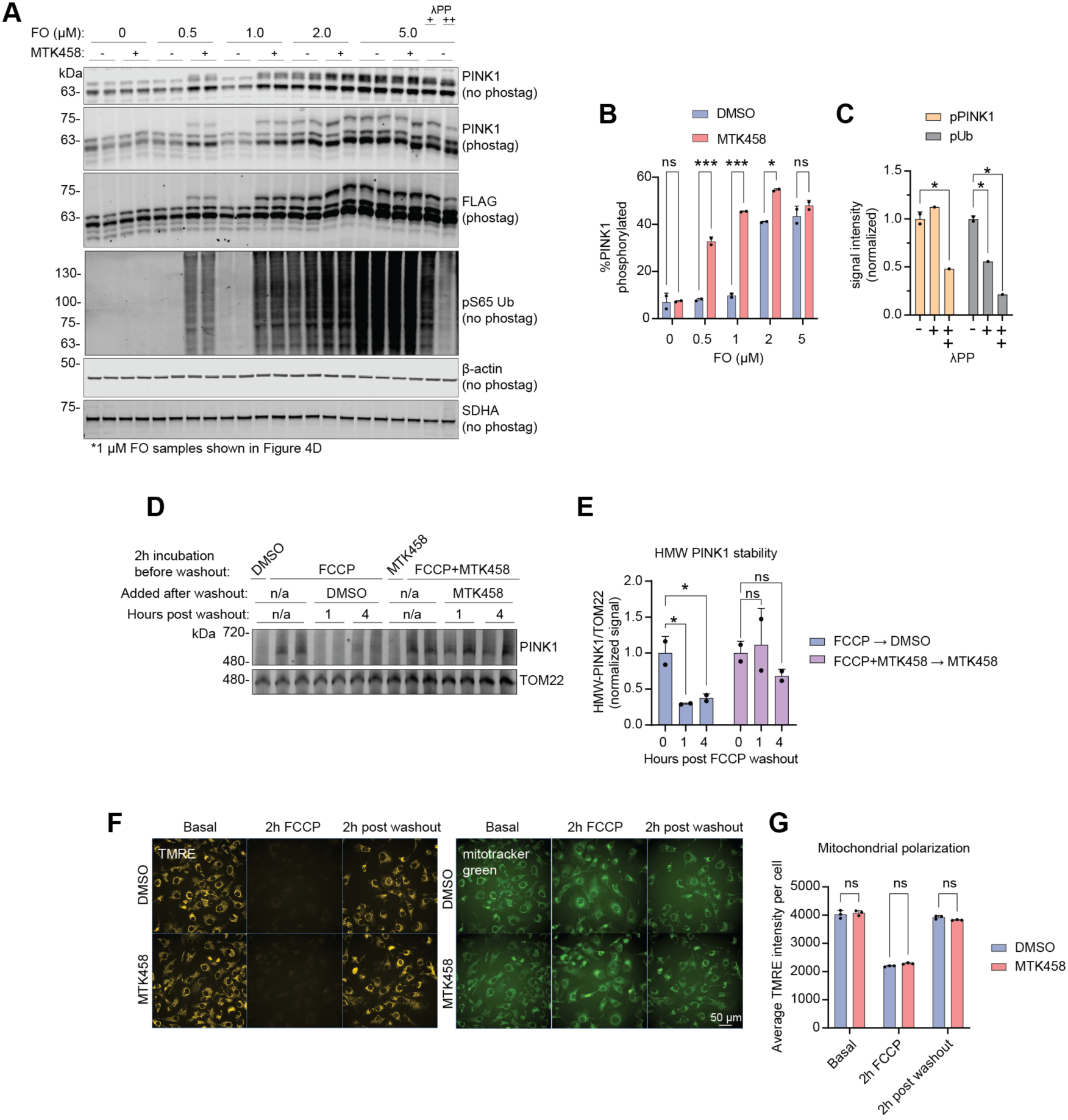
MTK458 stabilizes the active form of PINK1. (A) EPF1 cells (which overexpress PINK1-FLAG) were treated with FO and MTK458, and lysates were analyzed by phos-tag SDS-PAGE. We observed an increase in a phospho-PINK1 species by MTK458. Treatment of cell lysates with lambda phosphatase (ÅPP) decreases the phospho-PINK1 signal. (B-C) Quantification of band intensities in (A) is plotted. (D-G) FCCP washout experiments are performed, similar to the schematic in Figure 4I. (D) EPF1 cells were treated with 10 µM FCCP ± 2.8 µM MTK458 for 2h, followed by FCCP washout and treatment with DMSO or MTK458 for 1-4 hours. Cells were lysed for analysis on BN-PAGE, showing that MTK458 sustains the high molecular weight PINK1 complex even after FCCP removal. (E) Quantification of the gel in (D) is shown. (F) SK-OV-3 cells were treated with 10 µM FCCP ± 2.8 µM MTK458 for 2h, followed by FCCP washout and treatment with DMSO or MTK458 for 2 hours. At the indicated timepoints, mitochondrial membrane potential was quantified by live imaging with 10 nM TMRE. As a control, cells were also imaged with the mitotracker green dye at 100 nM. TMRE only enters polarized mitochondria, but mitotracker green enters all mitochondria. Scale bar, 50 µm. (G) Quantification of the TMRE intensity in (F) is shown. Mean ± SD. *p < 0.05, **p < 0.01, ***p < 0.001, ****p<0.0001, n.s., not significant.

**Supplementary Figure S6:**
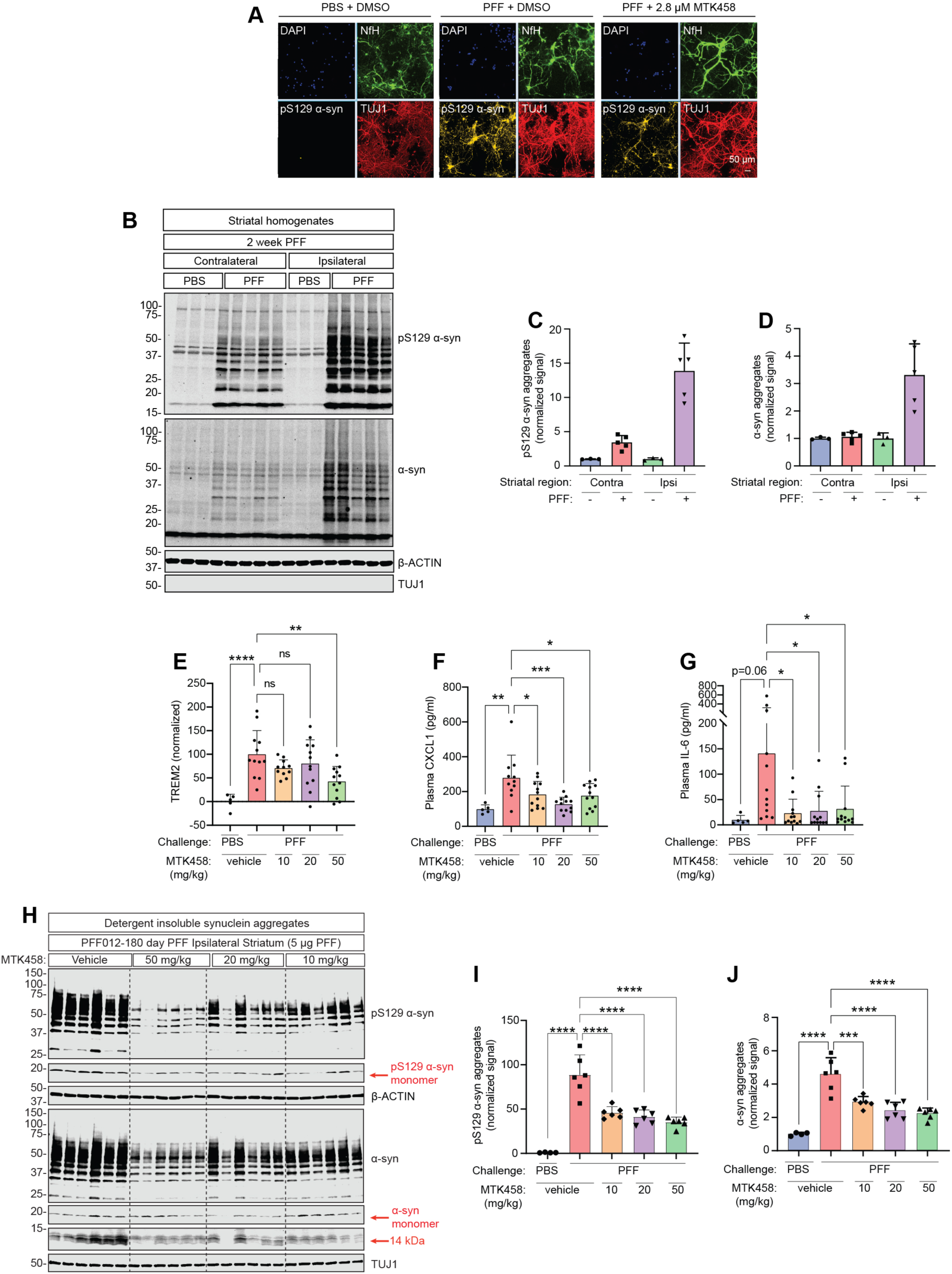
MTK458 decreases α-synuclein aggregates in vitro and in vivo. (A) Related to Figure 5F-I. Immunostaining shows representative images of neurons treated with mPFFs and 2.8 µM MTK458. Primary hippocampal neurons were seeded for immunostaining, treated with mPFFs (0.5 µg/mL) and MTK458 as in (E), and on DIV14 fixed for immunostaining with antibodies against pS129 α-syn, NfH and TUJ1. Scale bar, 50 µm. (B) Mice were challenged with striatal injection of PFFs of PBS. After 2 weeks, ipsilateral or contralateral striatum brain pieces were harvested for sequential extraction with buffers containing increasing amounts of detergent. NP-40 insoluble fractions were analyzed by immunoblot. (C-D) Quantification of (B) is shown. (E-G) Mice were challenged with striatal injection of PBS or PFF, and then dosed with MTK458 for 6 months. Mice were analyzed for TREM2 in the ipsilateral striatum (E), plasma CXCL1 (F), and plasma IL-6 (G) levels. (H) Similar experiment to Figure 6B, except that mice were sacrificed 180 days after PFF injection and MTK458 dosing (QD, PO) at the indicated concentrations. (I-J) Quantification of (H) is shown. Mean ± SD. *p < 0.05, **p < 0.01, ***p < 0.001, ****p<0.0001, n.s., not significant.

**Supplementary Figure S7:**
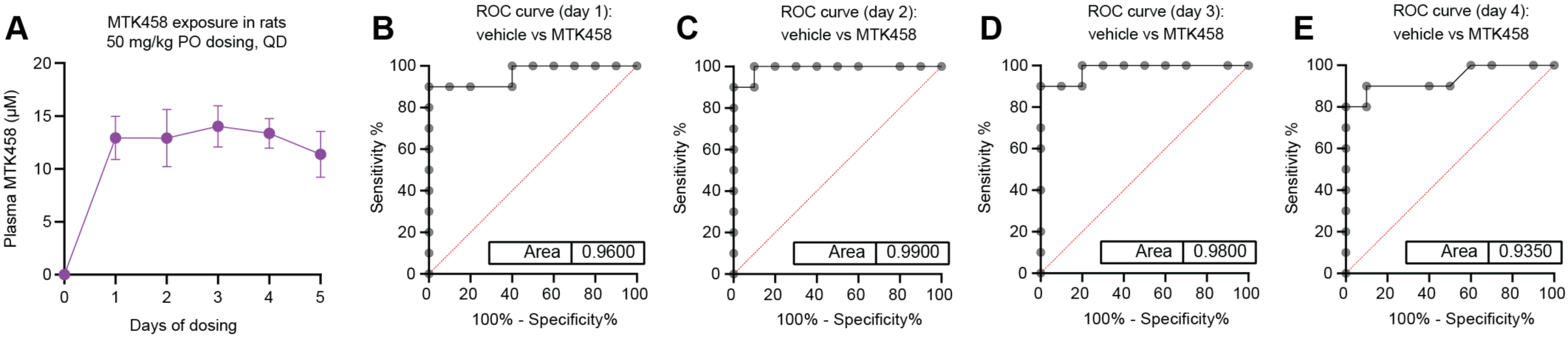
MTK458 dosing in naïve rats decreases plasma pUb. Related to Figure 6I-K. (A) Sprague Dawley rats were dosed (PO, QD) with 50 mg/kg MTK458 for 5 days (6 doses), and plasma was drawn before dosing, or 3h after every dose. Plasma MTK458 concentrations were determined by mass spectrometry and plotted. Mean ± SD is shown. (B-E) ROC curves for plasma pUb levels (normalized to predosing levels) in vehicle or MTK458 dosed rats (after 1, 2, 3, or 4 days of dosing) are shown.

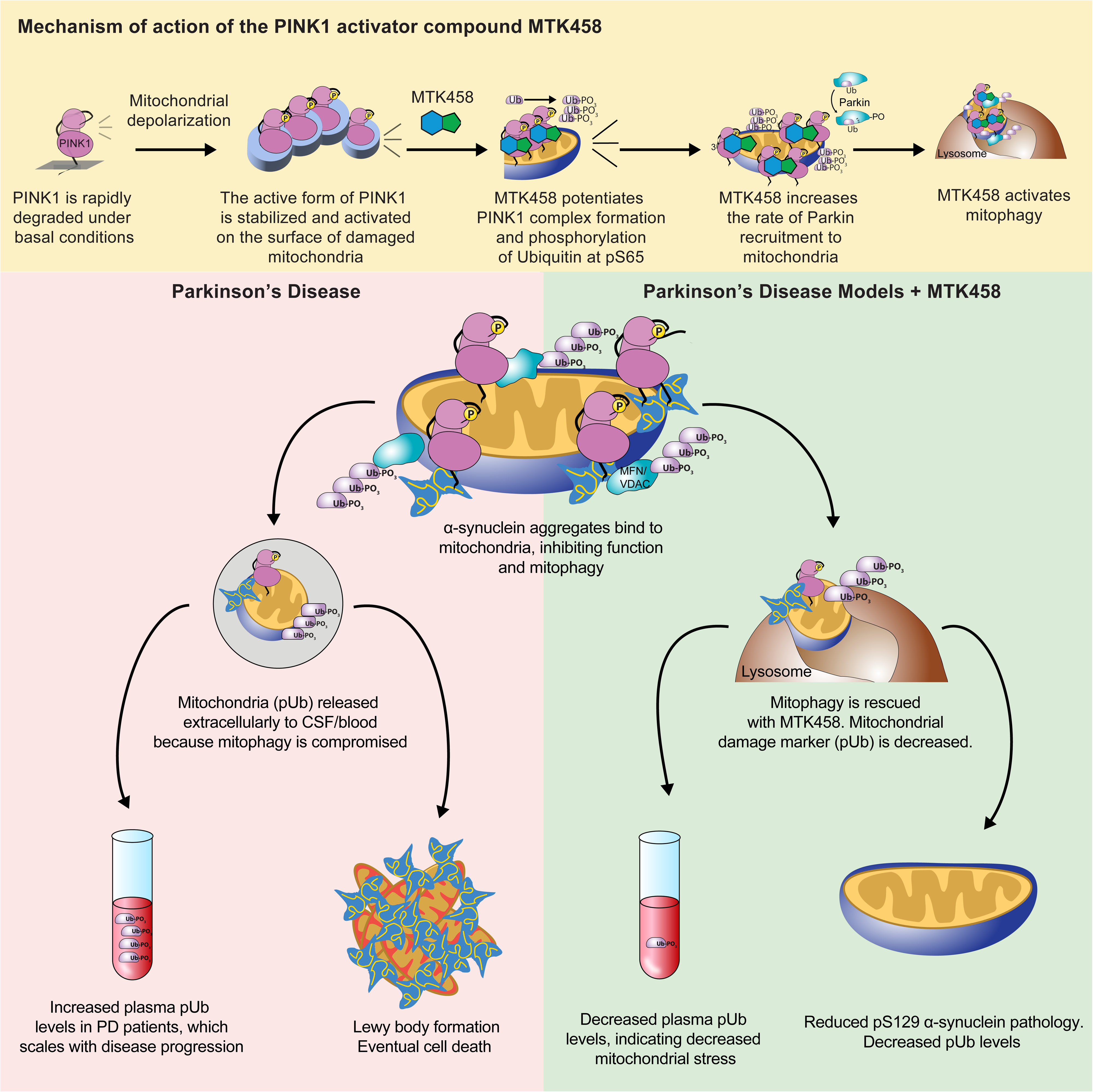

## KEY RESOURCES TABLE

### TABLE FOR AUTHOR TO COMPLETE

*Please upload the completed table as a separate document. **<u>Please do not add subheadings to the key resources table.</u>** If you wish to make an entry that does not fall into one of the subheadings below, please contact your handlin**g** editor. **<u>Any subheadings not relevant to your study can be skipped.</u>** (**NOTE:** For authors publishing in Cell Genomics, Cell Reports Medicine, Current Biology, and Med, please note that references within the KRT should be **in** numbered style rather than Harvard.)*

#### Key resources table

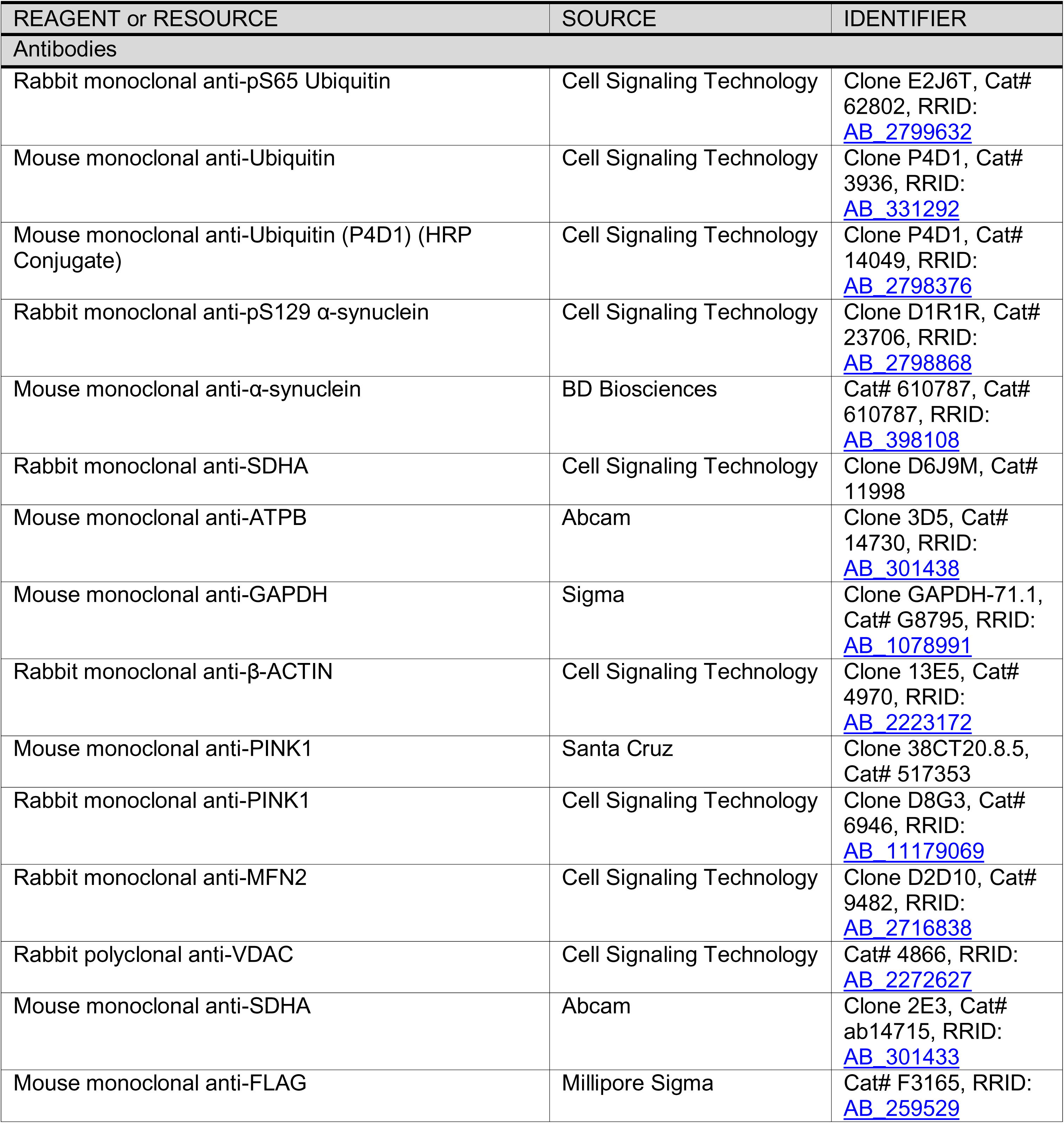
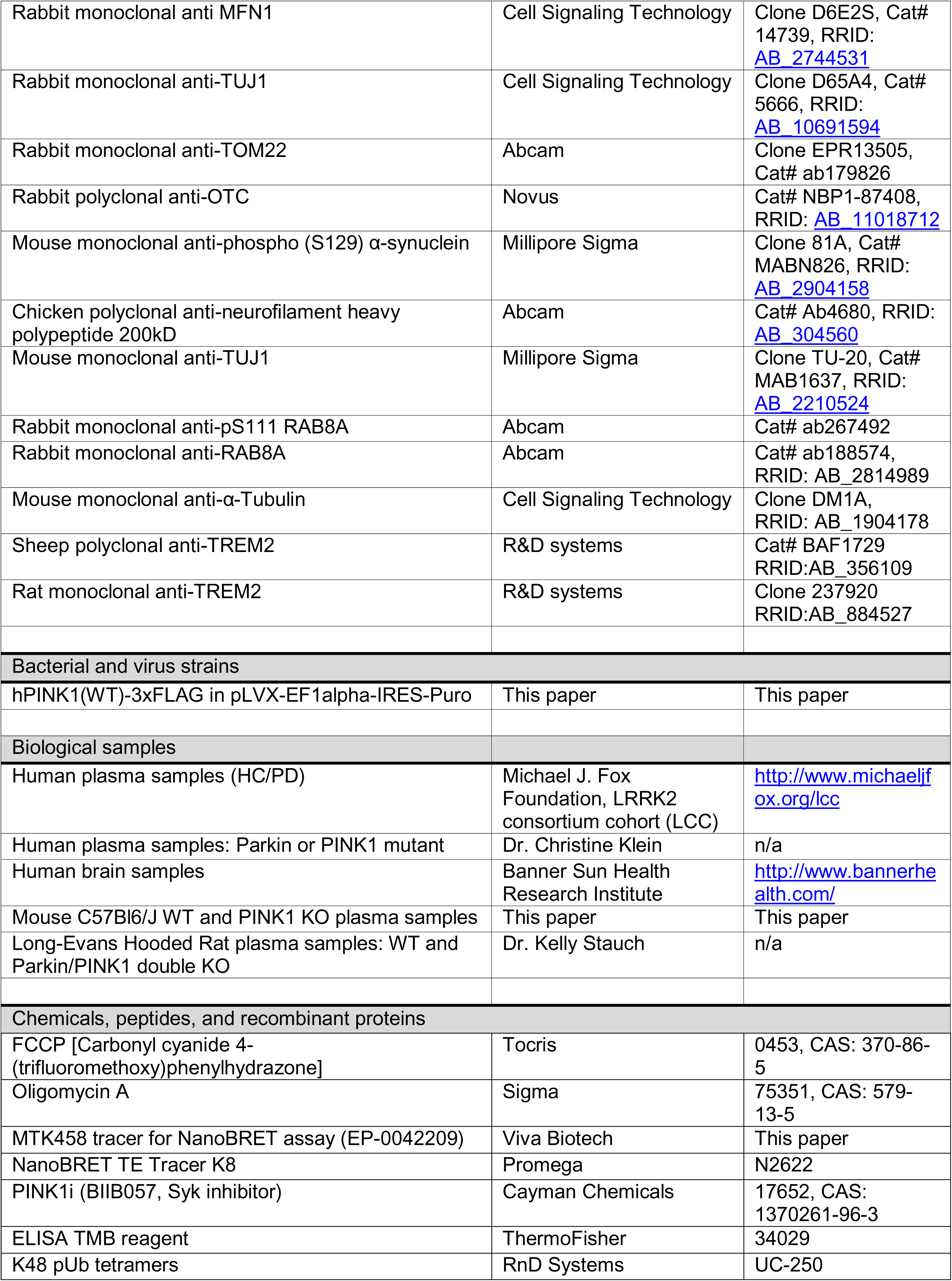
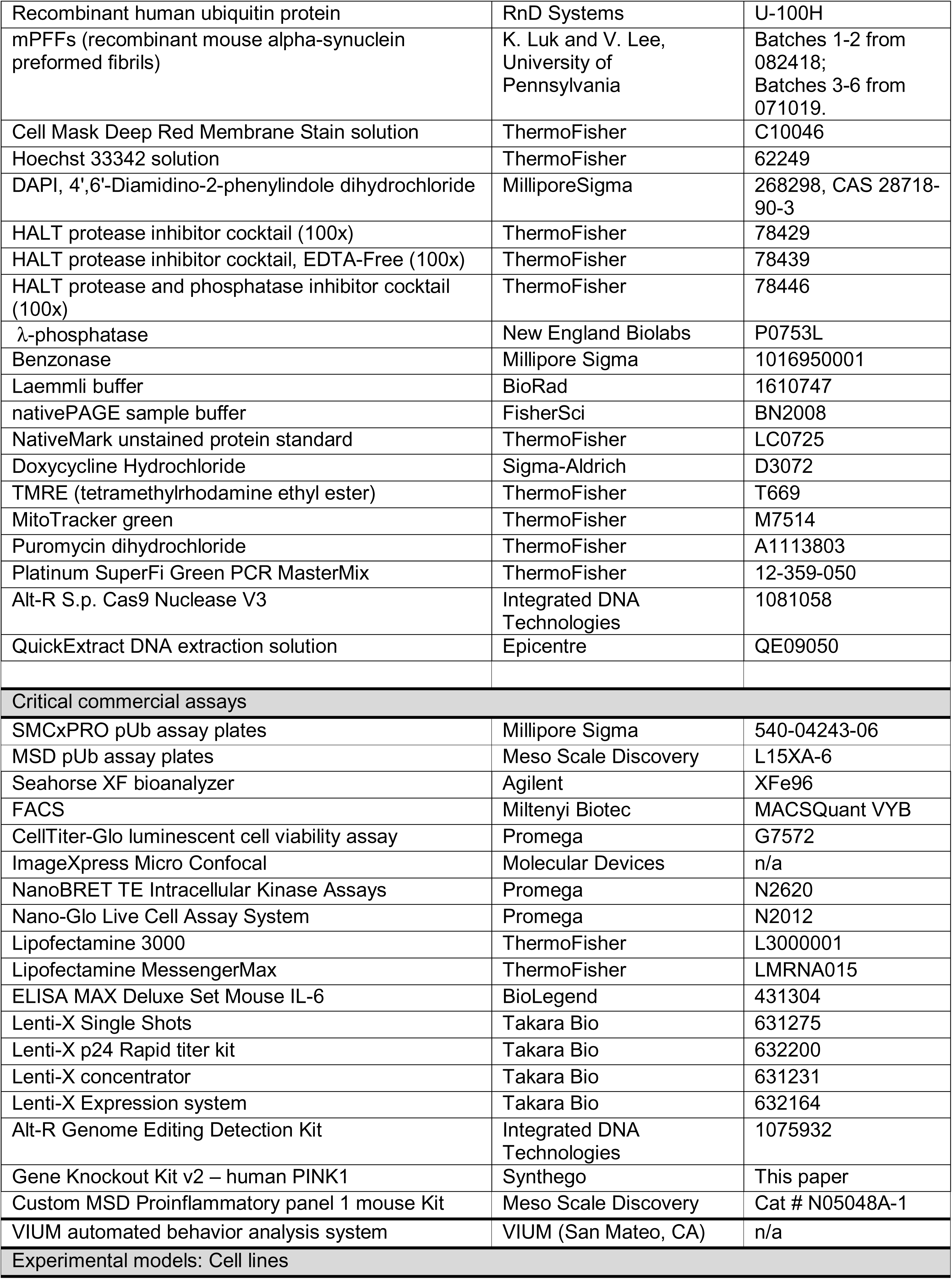
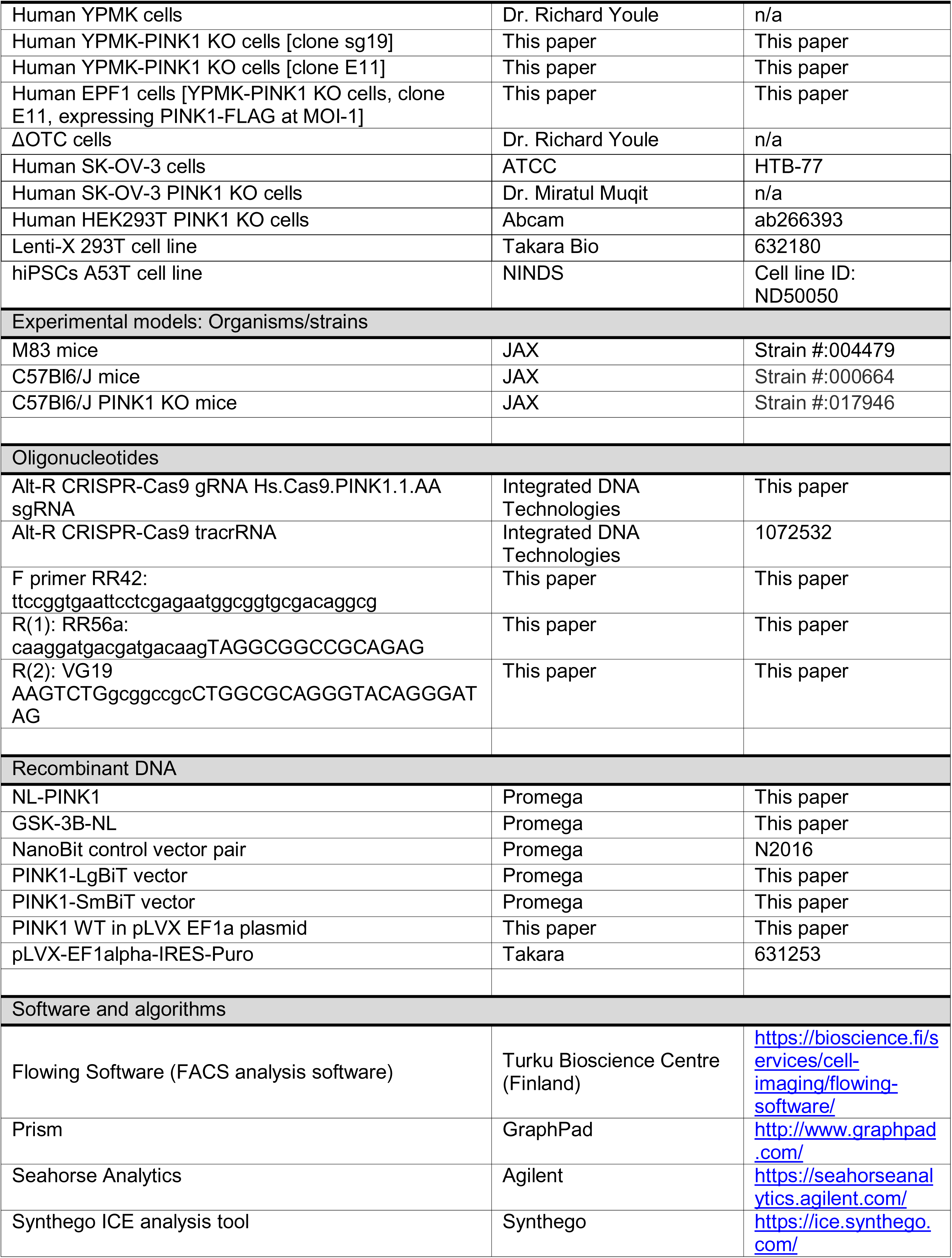
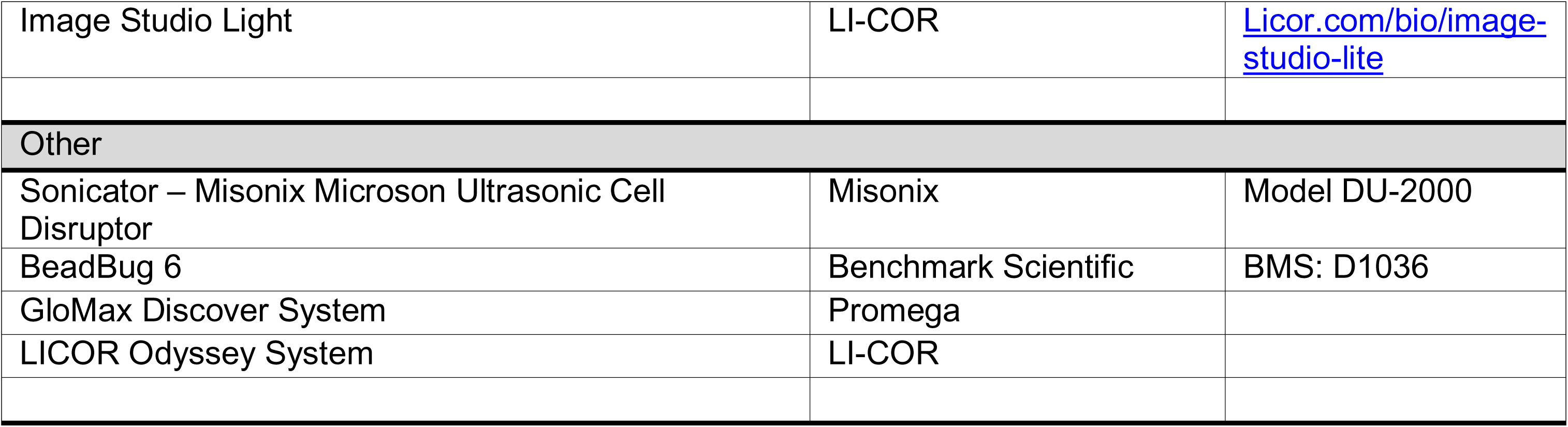

## STAR METHODS

### KEY RESOURCES TABLE

See Word document

## RESOURCE AVAILABILITY

### Lead contact

Further information and requests for resources and reagents should be directed to and will be fulfilled by the lead contact

### Material availability

Unique reagents generated in this study, such as cell lines, are available from the lead contact with a completed Material Transfer Agreement.

### Data and code availability

- All data and materials presented in this manuscript are available from the corresponding author upon a reasonable request under a completed Material Transfer Agreement.
- This paper does not report original code.
- Any additional information required to reanalyze the data reported in this work paper is available from the lead contact upon request.

## EXPERIMENTAL MODEL AND SUBJECT DETAILS

### Animals and ethics statement

Experiments were conducted with C57Bl6/J mice ordered from The Jackson Laboratories (Bar Harbor, Maine). All animal experiments were reviewed and approved by IACUC and done in accordance with the Animal Welfare Act (9 CFR, Part 1, 2 and 3) and the Guide for the Care and Use of Laboratory Animals guidelines (8th Edition). Mice were caged in Innovive Innorack Caging System and had ad libitum access to water and Global 18% Protein Rodent Diet from Envigo. During the study, the care and use of animals were conducted in accordance with the regulations of the Association for Assessment and Accreditation of Laboratory Animal Care (AAALAC).

Experiments with Sprague Dawley rats were done at BioDuro-Sundia (Shanghai, China). All animal experiments were reviewed and approved by IACUC and done in accordance with the Animal Welfare Act, the Guide for the Care and Use of Laboratory Animals guidelines. Rats were ordered from Zhejiang Vital River Laboratory Animal Technology Co.,Ltd, caged in Static Caging System fashion (3 rat/cage), and fed experimental rat maintenance feed (Shanghai Protein Bio-Technology Limited, Shanghai, China) diet. During the study, the care and use of animals will be conducted in accordance with the regulations of the Association for Assessment and Accreditation of Laboratory Animal Care (AAALAC, accreditation number is 001516).

### Human plasma samples

Biospecimens used in the analyses presented in this article were obtained from the MJFF-sponsored LRRK2 Cohort Consortium (LCC). For up-to-date information on the study, visit www.michaeljfox.org/lcc). Blood was collected from patients into K_3_EDTA tubes, inverted 8-10 times to mix, and centrifuged at 4°C at 1500 x g for 15 minutes (within 30 minutes of blood draw). Plasma was aliquoted at stored at −80°C. Samples were de-identified. Samples were analyzed in a blinded fashion on the SMCxPRO pUb assay. After all SMCxPRO runs were completed, samples were unblinded with respect to PD status, gender, and LRRK2 mutation status. UPDRS Part 3 total score (UPDRS3), Modified H&Y score, age, levodopa dosage, cigarette smoking habits, and/or coffee drinking habits were provided for some, but not all, of patients. A healthy control (HC) patient is defined as someone without PD.

### SMCxPRO pUb assay

SMCxPRO pUb assay was performed according to the manufacturer’s standard protocols. As a capture antibody, rabbit anti-pS65-Ub antibody (CST E2J6T lot 5) was coated onto SMC magnetic beads using the Capture Antibody labeling kit (Millipore Sigma 03-0077-02). As a detection antibody, mouse anti-Ub antibody (CST P4D1 lot 15) was labeled with Alexa Fluor dyes using the Detection Antibody labeling kit (Millipore Sigma 03-0076-02). For the calibration curve, K48 pUb tetramers (RnD systems UC-250) were used. To determine the concentration of the K48 pUb tetramers, a BCA assay using Recombinant Human Ubiquitin Protein (RnD systems U-100H) as the calibrator was performed. Assay plates (Millipore Sigma 540-04243-06) were read on the SMCxPRO system (Millipore Sigma 95-0100-00).

### Preformed Fibrils of α-synuclein (PFFs)

For PFF experiments for seahorse or mKeima analysis: wildtype full-length human α-synuclein PFFs were generated as described previously (Polinski et al., 2018) and obtained from the Luk lab at the University of Pennsylvania. PFFs were diluted to 0.1mg/ml in Dulbecco’s PBS (DPBS) and sonicated for 1 min at 30% amplitude with 1s on, 1s off pulses using a QSonica water bath sonicator.

For all other experiments: recombinant mouse WT α-synuclein pre-formed fibrils (mPFFs) were generously provided by K. Luk and V. Lee (UPenn, under MTA) at 5mg/mL. Prior to performing experiments with a new batch, a quality control assay was performed to compare the seeding efficiency between new and previous batch of mPFFs.

### Primary neuronal culture for PFF seahorse and mitophagy

Primary neuronal cultures were generated by dissection of hippocampal and cerebral cortex tissues from embryonic day 16 (E16) from A53T α-synuclein transgenic M83 mice according to Abbvie’s IACUC approved protocols. Briefly, tissues were dissected in ice-cold HBSS (Invitrogen, Cat# 14175095), separated from the meninges, and washed 3x in HBSS on ice. Tissues were then dissociated by incubation with neuronal isolation enzyme (ThermoFisher Scientific, Cat# 88280) at 30°C for 30min; the tubes were gently inverted every 5min to enable efficient enzyme digestion. After 30min, the tissues were gently triturated by manual pipetting 5-10x, resuspended in 8ml of plating medium (DMEM [Invitrogen, Cat# 11960044] supplemented with 10% fetal bovine serum, 1x penicillin-streptomycin), and filtered using a 70µm cell strainer (Falcon, Cat# 087712). Cells were then counted and plated in 150ul plating medium per well in poly-D-lysine pre-coated 96-well plates (Corning, Cat# 353219). 4h later, plating medium was aspirated and replaced with neuron maintenance medium (Neurobasal Plus [Invitrogen, Cat# A3582901] supplemented with B-27 Plus [Invitrogen, Cat# A3582801], 1x penicillin-streptomycin, and 1x GlutaMAX [Invitrogen, Cat# 35050061]). 1/3 volume fresh neuronal maintenance medium exchange was performed twice weekly until day of analysis.

### Seahorse respirometry in primary neurons

For PFF concentration curve: sonicated PFFs were added on DIV7, and cells were analyzed by Seahorse on DIV28. For PFF timecourse: sonicated PFFs were added on DIV7, DIV14, DIV21, or DIV28, and cells were analyzed by Seahorse on DIV35. Mitochondrial oxygen consumption rates (OCR) were measured using Agilent Seahorse XFe96 analyzer. Mouse primary neurons (30,000 per well) were plated in a 96-well XF Cell Culture Microplate. On the day of analysis, mitochondrial OCR’s were measured using XF Cell Mito Stress Test kit (Agilent, 103015-100), following the manufacturer’s instructions. Briefly, on the day of analysis, cells were washed 2× with and incubated in Mito Stress Test Assay Medium (1 mM pyruvate, 2 mM glutamine, and 10 mM glucose in XF base medium) in a 37 °C non-CO2 incubator for 1 h. OCRs were measured with an Agilent Seahorse XFe96 analyzer at baseline, after addition of 1.5 µM oligomycin to evaluate respiration associated with cellular ATP production, after addition of 1 µM FCCP to evaluate uncoupled respiration, and after addition of 0.5 µM antimycin/rotenone to measure non mitochondrial respiration. OCR were then normalized to the number of cells counted using brightfield imaging employing ImageXpress Micro Confocal High-Content Imaging System.

### Mitophagy in primary neurons

To measure mitophagy, primary neurons were treated with lentivirus expressing MT-mKeima-Red Fluorescent protein (MBL International Corporation) on day-in-vitro 4 (DIV4, 5 MOI). On day 7, the neurons were treated with Human PFF (1 µg/ml). The fluorescent protein Keima has an excitation spectrum that changes according to pH. A short wavelength (440 nm) is predominant for excitation in a neutral environment, whereas a long wavelength (586 nm) is predominant in an acidic environment. The ratio of fluorescent intensity in each excitation condition is an indicator of mitophagy in living cells. Mitophagy rate (586/440 ratio) was measured in live cells 21 days after PFF treatment using ImageXpress Micro Confocal (Molecular Devices).

### Cell lines and primary cultures

YPMK, YPMK PINK1 KO, EPF1, HEK293 PINK1 KO, SK-OV-3, and ΔOTC cell lines were maintained in our laboratory. YPMK cells were obtained from Dr. Richard Youle. SK-OV-3 cells were obtained from ATCC (HTB-77). YPMK PINK1 KO cells were a stable clone generated in our laboratory using CRISPR as described below. EPF1 cells are YPMK PINK1 KO cells that were infected with lentivirus to constitutively express PINK1 with a 3x FLAG tag at the C-terminal end. ΔOTC cells were kindly provided by Dr. Richard Youle. ΔOTC cells are HeLa cells stably expressing YFP-Parkin and doxycycline-inducible ΔOTC. These cell lines were maintained by culturing in DMEM (Corning 10-013-CV) with 10% FBS (Corning 35-010-CV) and penicillin-streptomycin (Corning 30-002-CI) in 5% CO_2_ incubator at 37°C.

### Sandwich pUb ELISA for YPMK cells

YPMK cells, maintained in culture medium [DMEM (Corning 10-013-CV) with 10% FBS (Corning 35-010-CV) and penicillin-streptomycin (Corning 30-002-CI)], were seeded in 96 well plates (Corning 3596) with test compounds at a concentration of 25 µM to 0.3 µM, at 10,000 cells/well. DMEM (Corning 10-013-CV) with 10% FBS (Corning 35-010-CV) and penicillin-streptomycin (Corning 30-002-CI) was used as the culturing medium throughout the assay. On the next day, medium was replaced with fresh medium containing 1 µM FCCP (Tocris 0453) and oligomycin (Sigma 75351) (FO) and test compounds. After 2h, cells were washed and 50 µl NH lysis buffer was added directly to the cells to lyse the cells. 10 µl of cell lysate was used in the sandwich pUb ELISA.

For the sandwich ELISA, rabbit anti-pS65 Ubiquitin antibody (CST, Clone E2J6T) was coated onto 96 well plates (Corning 3690) as the capture antibody. Mouse anti-Ubiquitin (HRP Conjugate, CST #14049) was used as the detection antibody. ThermoFisher (34029) 1-step ELISA TMB reagent was used to quantify the pUb signal per well. Signal intensities were normalized to control wells and graphed.

### mKeima assay

YPMK cells, maintained in culture medium [DMEM (Corning 10-013-CV) with 10% FBS (Corning 35-010-CV) and penicillin-streptomycin (Corning 30-002-CI)], were seeded in 96 well plates (Corning 3596) with test compounds at a concentration of 25 µM to 0.3 µM, at 10,000 cells/well. DMEM (Corning 10-013-CV) with 10% FBS (Corning 35-010-CV) and penicillin-streptomycin (Corning 30-002-CI) was used as the culturing medium throughout the assay. On the next day, medium was replaced with fresh medium containing 1 µM FCCP (Tocris 0453) and oligomycin (Sigma 75351) (FO) and test compounds. After 6h, cells were detached from the wells using trypsin, washed, and then resuspended in sorting buffer (145 mM NaCl, 5 mM KCl, 1.8 mM CaCl_2_, 0.8 mM MgCl_2_, 10 mM HEPES, 10 mM glucose, 0.1% BSA, 0.5 µg/mL DAPI). Cells were analyzed using FACS (MACSQuant VYB) with appropriate filters. For quantifying the acidic mKeima (localized in lysosomes), 561/620 nm excitation/emission filters were used. For quantifying the neutral mKeima (localized in mitochondria), 488/614 nm excitation/emission filters were used. The ratio of lysosomal to mitochondrial mKeima per cell indicates whether the cell is undergoing mitophagy (Supplementary Fig. 1A). The percentage of cells undergoing mitophagy under all concentrations of test compounds are graphed, and EC50 values from these curves were determined by Prism (GraphPad).

### Galactose/glucose assay

SK-OV-3 cells were maintained in culture medium [DMEM (Corning 10-013-CV) with 10% FBS (Corning 35-010-CV) and penicillin-streptomycin (Corning 30-002-CI)]. Prior to the assay, cells were trypsinized, washed three times in either glucose or galactose medium, and then seeded in 96 well plates (Corning 3610) at 9,000 cells/well. Glucose medium: DMEM with glucose (Corning 10-013-CV) + 10% FBS + 1x Penicillin-Streptomycin. Galactose medium: DMEM without glucose (Gibco 11966-025) + 10% FBS + 1x Penicillin-Streptomycin +1 mM pyruvate (Lonza 13-115E) + 10 mM galactose (Fisher G1-100). At 23-24h after cell seeding, medium was exchanged for fresh medium containing test compounds at 50 µM to 0.3 µM. At 19-20h after addition of test compounds, medium was removed and replaced with 50 µl glucose medium per well. Immediately after, the CellTiter-Glo assay (Promega G7572) was performed according to manufacturer’s instructions to assess cell number after test compound treatment. Luminescence was measured using the Promega GloMax Discover Microplate Reader. Within each medium, the number of cells remaining after test compound treatment was normalized to cells treated with DMSO alone. Then, the ratio of cell survival in galactose medium vs glucose medium was plotted for each concentration of test compounds. EC20 values from these curves were determined by Prism (GraphPad).

### Mitochondrial respiration analysis (Seahorse) in HeLa cells

HeLa cells (P4-7 after receiving from ATCC) were maintained in culture medium [DMEM (Corning 10-013-CV) with 10% FBS (Corning 35-010-CV) and penicillin-streptomycin (Corning 30-002-CI)], and plated into Seahorse XFe96 flux plates 24h before the assay. Seahorse XFe96 respirometry was performed in non-fluorescent DMEM, buffered with 10mM TES, 1mM NaHCO3, containing 25mM glucose, 1mM pyruvate, 2mM glutamine and 10% FBS. Compounds were added in addition port A at final concentrations of 0.1, 0.5, 2.5, 10 and 25 µM, with a final DMSO concentration of 1:2000, constant for all conditions. The experiment was performed in triplicate. The effects of the compounds on oxygen consumption rate (OCR) were monitored for one hour. This was followed by the standard mitochondrial stress test by first adding (port B) oligomycin at 2 µg/ml final concentration, then FCCP at 2 µM (port C; titrated here to elicit a maximal response), and finally antimycin A plus myxothiazol (2 µM each; port D). Basal respiration = [OCR 1h after MTK458 injection] – [non-mitochondrial respiration]. Maximal respiration = [OCR after FCCP injection] - [non-mitochondrial respiration]. Spare respiratory capacity = [maximal respiration] – [basal respiration].

### Generation of cell lines by CRISPR/Cas9 engineering (for PINK1-knockout), or lentiviral infection (stably expressing PINK1-3xFLAG)

#### Transfection with sgRNA-Cas9 RNP complex

Clonal PINK1-knockout cell lines were generated in parental HeLa YPMK cells (expressing YFP-Parkin and mt-mKeima) using either the Alt-R CRISPR-Cas9 System from IDT, or the Synthego 2.0 Gene Knockout kit, according to the manufacturer’s protocols. The sgRNA from IDT targeted exon 1 in *PINK1*, while the kit from Synthego included 3 sets of guide RNAs. In both approaches, guide RNAs targeting PINK1 along with purified Cas9-NLS. sgRNA:Cas9 RNP complexes were assembled at a ratio of either 1:1 or 1.3:1 (respectively), for a final concentration of either 10 µM or 20 µM, and then reverse transfected using Lipofectamine MessengerMax, seeding 5k cells / well in 96-well plates. At 48 h cells were passaged and DNA was extracted from an aliquot of cells using QuickExtract Solution. DNA from the bulk transfected pool was sequenced using primers around the relevant exon to confirm disruption in PINK1 sequence.

Upon selection of optimal conditions, cells from the master plate were passaged to identify single cell clones by limited dilution, plating 30 cells / 96-well plate. At 2 days post-seeding, 96-well plates were imaged by high content microscopy at 4x (Molecular Devices), to score positive wells and exclude any wells with more than one cell or colony. At ∼2 weeks, wells with single colonies were harvested, and 60 clones / plate were consolidated to maintain a master plate. Following DNA extraction, DNA from was submitted for sequencing analysis (GeneWiz). Sequences were analyzed using the Synthego ICE Analysis tool to identify clones with good knockout scores and sizeable deletion fragments. Multiple clones with high knockout scores were further evaluated in functional assays to confirm 1) the absence of PINK1 protein expression and lack of phosphorylated of ubiquitin, 2) lack of mitophagy induction, and 3) absence of parkin recruitment under conditions of high FCCP / Oligomycin treatment (by western blotting from isolated mitochondria, mKeima FACS assay, or live cell imaging, respectively). The best performing clones were used in subsequent experiments.

Following an initial round of cell line generation using only a single gRNA approach (IDT), even the best clones still showed some response to high F/O conditions at late timepoints, despite no detectable PINK1 protein or pS65 Ub signal. Clones arising from the multi-guide sgRNA approach (Synthego) overall had higher KO scores (>90), and larger deletions (>100nt), with no induction of mitophagy observed after high F/O conditions. From this multi-guide approach, one of the best clones, E11, was chosen for other assays.

### Infection with hPINK1(WT)-3xFLAG lentivirus

A lentivirus infection protocol was optimized to obtain expression levels of 3xFLAG-tagged PINK1 matching endogenous expression in the PINK1 in the HeLa YPMK E11 (PINK1 KO) background. lentivirus was made from 293T cells transfected with hPINK1(WT)-3xFLAG plasmid (a C-terminal tag), using the Lenti-X packaging Single Shots protocol. Following quantification of viral titer using a p24 ELISA kit, varying amounts of virus were used to infect cells at a range of MOI levels (in medium containing 8 µg/mL polybrene). Two days after infection, medium was replaced with medium containing 1–3 µg/mL Puromycin. One to two weeks following puromycin selection, stable pools were assayed for PINK1 expression level and pS65 Ub, or induction of mitophagy in the presence of high concentrations of F/O (by western blotting from mitochondrial fractions, or by mKeima FACS assay, respectively). From this approach, MOI-1 was selected as the optimal condition, and a stable cell line termed EPF1 (E11+ PINK1-3xFLAG at MOI-1) was used for other experiments (blue native gels).

### Human brain samples

De-identified human brain samples from HC or PD patients were purchased from Banner Sun Health Research Institute (Sun City, AZ). Brain samples were from the temporal lobe cortex region, and were stored frozen at −80°C until use. Brain samples were wrapped in aluminum foil and pulverized by hammer on a liquid nitrogen chilled surface. Approximately half of the pulverized brain was used for whole cell lysate, and the other half was used for mitochondrial/cytoplasmic fractionation.

For whole cell lysate, pulverized brain was added to RIPA buffer (Thermo Fisher Cat#89900) and “Triple pure” zirconium beads (Homogenizers, Cat# D1132-15TP), and lysed using the BeadBug 6 (Benchmark Scientific Cat# BMS:D1036) bead beater at maximum speed for 10 seconds. Lysates were transferred to a new tube and further homogenized using sonication (4 strokes; Misonix Microson Ultrasonic Cell Disruptor Model No. DU-2000) on ice. Lysates were incubated on ice for 30 minutes, and then centrifuged at 18,000 x g for 20 minutes at 4°C to remove bulk tissue and cell debris. Supernatant (whole cell lysate) was transferred to a new tube and stored at −80°C until analysis.

For mitochondrial and cytoplasmic fractionation, see below.

### Mitochondrial isolation from tissue lysates or cells

Mitochondria was isolated from frozen human brain, mouse striatum brain chunks, or mouse kidneys using the following protocol. Tissues were placed into tubes containing zirconium beads (Homogenizers D1132-15TP) and mitochondrial isolation buffer (MIB: 50 mM Tris·HCl (pH 7.5), 70 mM Sucrose, 210 mM Sorbitol, 1 mM EDTA, 1 mM EGTA, 100 mM Chloroacetamide, 1x Halt Protease and Phosphatase Inhibitor Cocktail (Thermo Fisher # 78440), 10 µM PR619). Tissues were homogenized using the BeadBug 6 (Benchmark Scientific Cat# BMS:D1036), followed by sonication at the lowest power level for 4 strokes (Misonix Microson Ultrasonic Cell Disruptor Model No. DU-2000). Homogenized sample was centrifuged at 300 RCF for 3 minutes at 4°C to remove bulk tissue. The supernatant was centrifuged again at 1,400 RCF for 10 minutes at 4°C to remove unbroken cells and other debris. The supernatant was centrifuged once again at 10,000 RCF for 10 minutes at 4°C, pelleting the mitochondria. The supernatant from this high speed centrifugation contains the cytoplasmic fraction and was saved for further analysis. The mitochondrial pellet washed in MIB and then lysed in NH lysis buffer (100 mM Bicine, 0.27 M Sucrose, 1 mM EDTA, 1 mM EGTA, 5 mM Na4P2O7, 100 mM Tris, 1% Triton X-100, HALT protease/phosphatase inhibitor cocktail (Thermo Fisher # 78440)) for Western blotting or ELISA/MSD analysis.

For cultured cells, cells were treated and collected by trypsinization. Cell pellets were washed with cold PBS and resuspended in MIB. Cells were lysed via sonication as above, and the same differential centrifugation steps were taken to isolate the mitochondria.

### Sandwich pUb MSD assay for tissue lysates

For the sandwich MSD assay, rabbit anti-pS65 Ubiquitin antibody (CST, Clone E2J6T) was coated onto MSD MULTI-ARRAY 96 well plates (MesoScale Discovery L15XA-3) as the capture antibody. Mouse anti-Ubiquitin (CST, Clone P4D1 was used as the detection antibody. SULFO-TAG labeled goat anti-mouse antibody (Meso Scale Discovery R32AC-5) was used as the secondary antibody. A K48 pUb tetramer calibrator curve (R&D Systems UC-250) was ran on every plate, and absorbance values from tissue lysates were interpolated back to the calibrator curve to calculate the pmol pUb per gram of lysate in the tissues.

### Mitochondrial Extract Proteomics

Mitochondrially derived ubiquitylated proteins were analyzed by Ubiquitin AQUA proteomics as described (Ordureau et al., 2015, Ordureau et al., 2014). Briefly, mitochondrial extracts were lysed in lysis buffer (50 mM Tris·HCl (pH 7.5), 1 mM EDTA, 1 mM EGTA, 50 mM NaF, 5 mM sodium pyrophosphate, 10 mM sodium 2-glycerol 1-phosphate, 1 mM sodium orthovanadate, 0.27 M sucrose, 1% (vol/vol) Nonidet P-40, 1µg/mL leupeptin/aprotinin, 0.5 mM 4(2-aminoethyl)benzenesulfonyl flouride(AEBSF)) containing 25 mM chloroacetamide and mitochondrial extracts were sonicated four times for a total of 15 s at the lowest settings and clarified by centrifugation (16,000×g for 10 min at 4 °C), and protein concentrations were determined by the Bradford assay. Mitochondrial extracts (15 µg) were subjected to TCA precipitation and digested for 6 hours at 37 °C with trypsin (in 100 mM tetraethylammonium bromide, 0.1% Rapigest (Waters Corporation)). Digests were acidified with an equal volume of 5% (vol/vol) formic acid (FA) to a pH of ∼2 for 30 min, dried down, resuspended in 5% (vol/vol) FA.

### UB-AQUA Proteomics

UB-AQUA was performed largely as described previously but with several modifications (Ordureau et al., 2015, Ordureau et al., 2014). A collection of 21 heavy-labeled reference peptides (Ordureau et al., 2015, Ordureau et al., 2014), each containing a single ^13^C/^15^N-labeled amino acid, was produced at Cell Signaling Technologies and quantified by amino acid analysis. UB-AQUA peptides from working stocks [in 5% (vol/vol) FA] were diluted into the digested sample [in 5% (vol/vol) FA] to be analyzed to an optimal final concentration predetermined for individual peptide. Samples and AQUA peptides were oxidized with 0.1% hydrogen peroxide for 30 min, subjected to C18 StageTip and resuspended in 5% (vol/vol) FA. Replicate experiments were performed and analyzed sequentially by LC/MS on an Orbitrap Qe-HFX instrument coupled with a Famos Autosampler (LC Packings, San Fransisco, CA) and an Accela600 LC pump (Thermo-Fisher Scientific). Peptides were separated on a 100 µm inner diameter microcapillary column packed in house with ∼35 cm of Accucore150 resin (2.6 µm, 150 Å, ThermoFisher Scientific, San Jose, CA). The column was equilibrated with buffer A (3% ACN + 0.125% FA). Peptides were loaded onto the column in 100% buffer A. Separation and elution from the column were achieved using a 100-min 0–25% gradient of buffer B [100% (vol/vol) ACN + 0.125% FA]. The scan sequence began with FTMS^1^ spectra (resolution of 120,000; mass range 300-1000 m/z; automatic gain control (AGC) target 5×10^5^, max injection time of 100 ms). The second scan sequence consisted of a targeted-MS^2^ (tMS^2^) method were MS^2^ precursors of interest were isolated using the quadrupole and analyzed in the Orbitrap (FTMS^2^) with a 1.0 Th isolation window, 30k resolution, 1×10^5^ AGC target and a max injection time of 200 ms. MS2 precursors were fragmented by HCD at a normalized collision energy (NCE) of 25%. LC-MS data analysis was performed using Skyline software (MacLean et al., 2010) with manual validation of precursors and fragments. The results exported to Excel and Prism (GraphPad) for further analysis and plotting. Total UB was determined as the average of the total UB calculated for each individual locus, unless specified otherwise.

### Western blotting and immunofluorescence for primary neurons

Primary neuronal hippocampal cultures were prepared from WT C57/BL6 mice at E17.5. Dissected brain tissue was collected into cold Hibernate E buffer, and then dissociated in papain solution containing DNAse for 30-45 min at 37°C. Following removal of papain solution and rinsing in plating medium (Neurobasal containing 10% FBS, B27, Glutamax, and P/S), tissue was gently triturated to disrupt tissue aggregates and then filtered through a 40μM strainer. Cells were seeded at a density of 300k cells / well in poly-D-lysine coated 24-well plates for Western blot studies, or XXX cells/well in poly-D-lysine coated 96-well plates for imaging studies. At 3h post plating, the plating medium was completely replaced with neuronal medium (Neurobasal, B27, Glutamax, and P/S). On DIV2 and DIV5, half the neuronal medium was replaced. On DIV7, mPFFs were sonicated and added at 0.5 µg/mL (according to Volpicelli et al 2014). On DIV9 and DIV12, medium was completely removed and replaced with medium containing MTK458 at various doses or DMSO. On DIV14, neurons were either collected for biochemical analysis (Western blotting) or used in immunofluorescence experiments. Immunofluorescence procedure is the same as in the ΔOTC cells. Cells were imaged on the ImageXpress Micro Confocal (Molecular Devices).

### Serial extraction of o-synuclein species from primary neurons, iPSC-neurons, or mouse brain tissue

For serial extraction, cells or tissue were collected in NP-40 lysis buffer (containing 10 mM Tris-HCl, pH 7.4, 150 mM NaCl, 5 mM EDTA, 0.5% Nonidet P-40, 1x Halt™ Protease and Phosphatase Inhibitor Cocktail, with Benzonase (1:1000). Following differential centrifugation at 22,000g for 20 min at 4°C, the NP-40 insoluble pellet was resuspended in SDS-Brain lysis buffer (containing 10 mM Tris-HCl, pH 7.4, 150 mM NaCl, 5 mM EDTA, 0.5% Nonidet P-40, 1% SDS, 0.5% sodium deoxycholate, 1x Halt™ Protease and Phosphatase Inhibitor Cocktail, with Benzonase (1:1000) and sonicated. Protein concentration was determined by BCA assay, and samples were analyzed for pS129 α-synuclein and total α-synuclein levels by western blot.

### Patient derived iPSC neuron assay

Human iPSCs were purchased from NINDS (https://stemcells.nindsgenetics.org/), and the hiPSCs cell line (Cell line ID: ND50050) with A53T mutation was used in this study. Human iPSCs were cultured using standard protocol with the complete mTeSR-1 medium (STEMCELL Technologies, #85851-basal medium, #85852-supplement) on Matrigel-coated plates (Corning, #353046). Human iPSCs were differentiated into DA neurons. Briefly, iPSCs were cultured on Matrigel (Corning)-coated plate in KnockOut™ Serum Replacement (Invitrogen, #10829-018, #10828-028) medium containing growth factors and small molecules including FGF8b (100 ng/mL, R&D Systems, #423-F8), SHH C24 (100 ng/mL, R&D Systems, #1845-SH), LDN193189 (100 nM, Stemgent, #04-0074), SB431542 (10 mM, Stemgent, #04-0010-10), CHIR99021 (3 µM, Stemgent, #04-0004), and purmorphamine (2 µM, Sigma, #540220) for the first five days. Afterwards, cells were maintained in neurobasal medium (ThermoFisher, #21103049) containing B-27 supplement (ThermoFisher, 17504-044), N-2 supplement (ThermoFisher, 17502-048) along with LDN193189 and CHIR99021 for six days. In the final stage, the cells were dissociated into single cell suspension and seeded on poly-ornithine and laminin coated plate in neurobasal medium containing B27 supplement, BDNF (20 ng/mL, Miltenyi Biotec, #130-096-286), GDNF (20 ng/mL, Miltenyi Biotec, #130-098-449), TGFb (1 ng/mL, Miltenyi Biotec, #130-094-007), ascorbic acid (0.2 mM, Sigma, A4034), dibutyryl-cAMP (0.5 mM, Sigma, D0627) and DAPT (10 µM, Tocris, #2634) until maturation. DA neurons were cultured for > 60 differentiation days before measurements. For differentiated human DA neurons were treated with different doses of MTK458 for 10 days before harvesting. For the F/O treatment, the differentiated human DA neurons were treated with different doses of F/O for 24 hours before harvesting. The treated cells were harvested and processed for mitochondria isolation, and further analyzed with Western blot as described in previous sections.

### α-Synuclein PFF *in vivo assay*

Male C57Bl6/J mice (8 – 10 wks of age) were anesthetized with an intraperitoneal injection of ketamine hydrochloride (100 mg/kg), xylazine (10 mg/kg), and acepromazine (1 mg/kg), and stereotaxically injected with recombinant α-synuclein fibrils (5 µg per brain). Control C57 animals received sterile PBS. A single needle insertion (coordinates: +0.5 mm relative to bregma, 2.0 mm from midline) into the right forebrain was used to target the striatum located at a depth of 2.8 mm below the dura. Material was injected via Stoelting QSI auto-injector at a rate of 0.2 µl per min (2 µl total per site) with the needle in place for 5 min at each target after injection. After recovery from surgery, animals were monitored regularly. At takedown, mice were perfused with PBS, euthanized, and brains were collected. The ipsilateral striatum brain chunks were subjected to serial extraction of α-synuclein species

### Mouse behavior analysis (Vium)

Mice were acclimated to single housing in Vium (San Mateo, CA) transparent cages for 7 days prior to behavioral analysis. Behavior for different parameters, were recorded for 7 consecutive nights. Built-in behavioral modules from Vium automated analysis were output for each of the 7 consecutive nights and treated as an independent experimental replicate.

### Mouse plasma IL-6 and CXCL1 assay

MSD detection used a duplex custom kit (Custom MSD Proinflammatory panel 1 mouse Kit, cat. no. N05048A-1) for IL-6 and KC/GRO (CXCL1). The kit contained a 96-well MSD plate, sample diluents, read buffer, antibodies, and antibody solution diluent. IL-6 and CXCL1 were simultaneously detected following the kit instructions. Samples were diluted 1:1 with sample diluent for readouts. Briefly, the plate was washed, then standard and samples were added to incubate for 2hrs at RT. After, the plate was washed three times, detection antibodies were added to incubate for 2hrs at RT. The plate was washed three times, and reading buffer was added to be read on the MSD instrument (MSD MESO QuickPlex SQ120, Rockville MD). Results were analyzed using Prism (GraphPad).

### TREM2 assay

TREM2 concentration was quantified using a custom sandwich MSD assay. Sheep anti-TREM2 polyclonal antibody (R&D systems BAF1729) was coated onto Small Spot Streptavidin 96 well plates (MSD, L45SA-2) as the capture antibody. Monoclonal rat anti-TREM2 antibody (R&D Systems, Clone # 237920) antibody was used as the detection antibody. SULFO-TAG labeled goat anti-mouse antibody (Meso Scale Discovery R32AC-5) was used as the secondary antibody. A calibrator curve of recombinant mouse TREM2 Fc Chimera Protein (R&D Systems, 1729-T2-050) was included on every plate, and absorbance values from tissue lysates were interpolated back to the calibrator curve to quantify the TREM2 in the tissue lysates.

### Mouse brain mitochondrial pUb assay

C57BL6J wildtype mice were challenged with PFF injection into the right striatum. After 3 months, were mice dosed with MTK458 (50 mg/kg, QD, PO with NMP:Solutol:water 10:10:80 v:v vehicle) for the indicated times. After the last dose, mice were perfused with PBS, euthanized, and brains were collected. The ipsilateral striatum brain chunks were subjected to mitochondrial isolation and lysates were analyzed for pUb content with the MSD pUb assay.

### Mouse plasma pUb assay

C57BL6J wildtype mice were challenged with PFF injection into the right striatum. The PFF groups were dosed with MTK458 (50 mg/kg, QD, PO with NMP:Solutol:water 10:10:80 v:v vehicle) for 1, 2, or 3 weeks. Plasma was collected from mice and analyzed by the SMCxPRO pUb assay.

### Rat plasma pUb assay

Experiments with Sprague Dawley Rats (7-9 weeks old) were performed by BioDuro (Shanghai, China). Rats were dosed with MTK458 (50 mg/kg, QD, PO with NMP:Solutol:water 10:10:80 v:v vehicle) for 6 doses. Blood was drawn by JVC (Jugular Vein Cannulation) method into K2-EDTA tubes and within 30 minutes of blood draw, blood was centrifuged at 4C, 1500 x g for 15 minutes. Supernatant (plasma) was transferred into new tubes and stored at −80C immediately. Plasma was sent to Mitokinin upon completion of experiment. Plasma was analyzed for pUb content by the SMCxPRO pUb assay.

